# Tomato Response Evaluation through Fertilization and PGRs application Under Temperature Differentiation in late Winter

**DOI:** 10.1101/2023.08.04.552040

**Authors:** Joydeb Gomasta, Jahidul Hassan, Hasina Sultana, Yukio Ozaki, Saud Alamri, Alanoud T. Alfagham, Latifah A AL-Humaid

## Abstract

This study evaluated the exogenous application of PGRs substitute chemical fertilization without compromising the growth and yield of tomato in fluctuated day-night temperature and humidity stressed late winter. Two-factor experiment comprising chemical fertilizers at 100, 110, 90 and 80 % of recommended doses besides control and PGRs of GA_3_; NAA, 4-CPA and SA @ 50 ppm including control was conducted where treatments were assigned in triplicates. Results revealed no significant variation among the fertilizer doses (80% to 110% of recommendation) regarding growth and yield contributing traits while among the PGRs, GA_3_ @ 50 ppm produced maximum number of flower clusters plant^-1^ (16.85), flowers (8.80) and fruits (5.79) cluster^-1^, single fruit weight (67.83 g) and fruit yield (6.61 kg plant^-1^) of tomato that was statistically identical with the findings of SA. But significant reduction in yield was noted in NAA and 4-CPA (1.20 kg and 1.21 kg plant^-1^, respectively). Interestingly, GA_3_ and SA in combination with any doses of the studied fertilizers maximize the tomato morphological and reproductive traits while fertilizer plus NAA and 4-CPA interaction gave the inferior results. Further, correlation matrix and PCA findings revealed that five fertilizer doses have no distinctiveness whereas GA_3_ and SA has distinct position than other PGRs with the maximum dependent variables those were contributed positively in the total variations. The study findings suggested that 20% fertilizer requirement could be reduced with the substitution of GA_3_ and SA @ 50 ppm for successful cultivation of tomato in late winter having the extreme environmental issues.

## Introduction

Tomato (*Solanum lycopersicum* L.) is one of the most extensively used and cultivated vegetables in the globe. The nutritional importance of tomato can be largely explained by its content of various health-promoting compounds, including vitamins, carotenoids, and phenolic compounds (Li et al., 2018). These bioactive compounds have a wide range of physiological properties, including anti-inflammatory, anti-allergenic, antimicrobial, vasodilatory, antithrombotic, cardio-protective, and antioxidant effects (Braga et al., 2016). Tomatoes are also rich in carotenoids, representing the main source of lycopene in the human diet (Viuda-Martos et al., 2014). Tomatoes also have the naturally occurring antioxidants vitamin C and E (Khan et al., 2021) as well as large amounts of metabolites, such as sucrose, hexoses, citrate, malate and ascorbic acid (Li et al., 2018). But the recent phenomenon of global warming across the world has posed severe challenges to vegetable production including tomato. Among others, the challenges include increase in air temperature (AT), fluctuation in atmospheric humidity (RH) and intensity of solar radiation (Meena et al., 2014). In Bangladesh, winter occurs for a short period being no longer than three months and temperature rise drastically in the post winter months with a variation between high and low pick for more than 15 °C that experienced in the recent years (Khan et al., 2019). These extremities in temperature and humidity phenomena sometimes becomes severe at the late winter and pre-summer period for successful cultivation of tomato. The optimal growth temperature for tomato growth is 18.3 to 32.2 °C, and the relative humidity is 50% to 70% (Shamshiri et al., 2018), however above 35 °C the growth is slow, and at 40 °C the plants stop growing (Firon et al., 2012). Excessive temperature rise causes poor pollination and reduces fruit setting (Singh et al., 2015), plant dwarfing, senescence (Lin et al., 2018) and quality deterioration (Mulholland et al., 2003).

About 251.69 million tons of tomato has been produced from 6.16 million hectares of land across the globe (FAOSTAT, 2022) with the gross annual production of 4.16 lakh tons from an area of 28.53 thousand hectares in Bangladesh (BBS, 2022). Although, dietary shortage of vegetables is still evident in the country. Again, to meet the daily vegetable requirement of 235 g person^-1^ day^-1^ the growers are encouraged to increase the total production but the only thing farmers are used to practice for increasing the yield and production is the excessive and indiscriminate use of inorganic fertilizers neglecting the recommended guidelines (Ahmmed et al., 2018); thereby rendering the soil health in danger for future production. Such overuse of inorganic fertilizers has caused soil, air, and water pollutions through nutrient leaching, destruction of soil physical characteristics, accumulation of toxic chemicals in water bodies, and so on (Kakar et al., 2020), as well as causing severe environmental problems and loss of biodiversity (Mozumder & Berrens, 2007). The continuous and steady application of inorganic fertilizers leads plant tissues to frequently absorb and accumulate heavy metals, which consequently decreases the nutritional and grain quality of crops too (Abebe et al., 2022; Lolamo et al., 2023). Therefore, reduction of using agrochemicals especially the use of chemicals fertilizer is very much urgent in the country as any means not only to improve the soil health but also to get quality vegetable product.

Again, plants use several physiological adaptive mechanisms such as hormonal changes, cellular or molecular adaptive mechanisms to survive environmental implications (Yadav and Sharma, 2016). Plant growth regulators (PGRs) perform a significant role in plant developmental process and thus modulate plant replies to abiotic stresses along with normal growth and developmental processes. PGRs have been implicated in efficient utilization of nutrients and translocation of photo-assimilates (Siddiqui et al., 2016). Exogenous PGRs can help to manage balance of phytohormones and thereby they trigger the plants’ tolerance to different stress. More specific responses include alteration of C partitioning, greater root: shoot ratios, enhanced photosynthesis, altered nutrient uptake, improved water status and altered crop canopy (Meshram et al., 2022). In recent years, exogenous plant growth regulator (PGR) treatment has been used to effectively improve crop drought and heat tolerance and preserve yield under salinity and drought stress (Zhang et al., 2022). Thus, plant hormones are the key regulators of plant growth and developmental processes as well as crucial for biotic and abiotic stress response throughout their life cycle (Hassan and Miyajima, 2019). However, the comprehensive study on the PGRs application in response to the fertilization minimization for crop cultivation are still scare. Therefore, it has been hypothesized that application of certain PGRs like auxin, gibberellic acid (GA_3_) and others might ascertain excellent tomato production under adverse environmental conditions with low chemical fertilizer requirements. Considering these, the present research was designed to assess the responses of tomato plants to varied levels of fertilizers from 20 % less to 10 % excess than recommendation after the foliar application of plant growth regulators.

## Materials and Methods

### Study Area and Planting material

The field and laboratory work of experiment was carried out at the research field and laboratory of the Department of Horticulture, Bangabandhu Sheikh Mujibur Rahman Agricultural University (BSMRAU), Gazipur-1706, Bangladesh during November 2022 to April 2023. Geographically the site was in 24.0379 °N and 90.3996 °E and characterized as a mix of tropical and sub-tropical climate with hot dry summer, long humid monsoon and short and dry winter (Khan et al., 2023). Healthy, insect-pest and pathogen free seeds of tomato var. *BARI Tomato-14* were collected from Bangladesh Agricultural Research Institute (BARI), Gazipur, Bangladesh prior to initiation of the experiment.

### Crop Management

Pot cultivation was practiced under semi-protected environment. Grey colored plastic pots of 30 cm depth and 30 cm diameter were filled with planting media prepared by mixing well decomposed cowdung, vermicompost and garden soil in the ratio of 2:1:4 (v/v). Nutrient fertilizers were also added to the media as per treatment. Pretreated seeds of the variety “*BARI Tomato-14*” were soaked overnight and sown in a plastic tray on 12^th^November 2022. Seeds were germinated within five days; young tender seedlings were then transplanted to polybags (4 cm × 6 cm) to get strong, healthy seedlings for planting in the main pots. Uniform growth seedling at twenty five-day old were transplanted to the previously prepared pots on7^th^ December 2022. Two seedlings were transplanted in each pot and single plant was retained upon establishment. Stalking was provided on 20^th^ December 2022 to protect the crop from lodging. All other intercultural operations like weeding, watering, insect-pest management etc. were performed as per commercial guides (Azad et al., 2020).

### Experiment Design and Layout

The pot experiment was set in a factorial randomized complete block design (RCBD) with three replicates where three pots each containing single plant were considered as a replication under each treatment. Treatments consisted of five different doses of fertilizers including control for the factor one and in factor two four separate types of plant growth regulators (PGRs) namely Gibberellic acid (GA_3_), Naphthalene acetic acid (NAA), 4-Chlorophenoxy acetic acid (4-CPA) and Salicylic acid (SA) @ 50 ppm of each PGRs were used along with control or no PGR. All the pots with tomato plants were placed 60 cm apart from each other for convenient intercultural operations.

### Treatment Preparation and Application

Nutrient treatment was given as soil feeding of fertilizers. Recommended fertilizer doses for tomato was accrued from FRG (Fertilizer Recommendation Guide), 2018 (Ahmed et al., 2018) of Bangladesh. Besides control (no fertilizer), the four fertilizer doses for individual pots of each replication were 100 %, 110 %, 90 % and 80 % of FRG’2018 where 100 % of FRG’2018 was denoted as 12 g of urea, 10 g of TSP, 5 g of MoP, 3 g of Gypsum, 0.5 g of ZnSO_4_ and 0.5 g of Boric acid plant^-1^.For applying nutrients; full of TSP, Gypsum, ZnSO_4_ and Boric acid and 1/3^rd^of MoP was mixed with the planting media during pot preparation. Urea and rest of the MoP was applied as side dressing in three and two installments, respectively. Urea was applied at the base of the plants and mixed with the media at 10, 25 and 40 days after transplanting (DAT). While, the 2/3^rd^ of MoP was mixed with the pot soil at 25 and 40 DAT. Immediately after fertilization, light irrigation was applied every time.

On the other hand, the PGR treatment was applied as foliar spray at the vegetative growth stage of the plants for twice at 23 and 38 DAT i.e., two days before fertilizer application so that PGR mediated physiological and hormonal activities could influence the nutrient uptake from the soil. All the PGRs were applied at 50 ppm concentration. Two litre (2 L) solution of each of the PGRs was prepared to spray 45 plants under each treatment. For preparing 1 L solution, exactly 50 mg of PGR powder was weighed in a 50 mL beaker and 5 mL of Methanol (70 %) was added into it and agitated in a magnetic stirrer until completely dissolved. Then, it was poured in a 1 L volumetric flask and filled up to the mark by adding distilled water. The procedure was repeated to make the solution volume to 2 L. Plants were sprayed with the solutions as per treatments at the both sides of the leaf until runoff from the leaves was noticed. Control plants were applied with equal volume of distilled water only at the similar way adding 10 mL of methanol in 2 L water.

### Measurement of Growth Characteristics

Influence of fertilizers and PGRs on vegetative growth characters of tomato were assessed by measuring plant height (cm), base diameter (cm), number of branches and leaves plant^-1^ and canopy spread (cm) periodically at 20, 45 and 75 DAT. Further, number of leaflets leaf^-1^, internode length (cm), individual leaf area (cm^2^) and leaf greenness (SPAD value) were determined at fruiting stage. Individual leaf area (cm^2^) and SPAD value of five leaves leaving 5-6 leaves from top were measured by an electric area meter (LI 3000, USA) and portable chlorophyll meter (SPAD-502Plus; Konica Minolta, Japan), respectively. Individual leaf area was estimated as per Aurdal et al. (2022) where leaflets of the large tomato leaf were separated and differently run under the electric area meter and averaged. Again, after the final fruit harvest at 29^th^ March, plants were carefully removed out of the pots, root systems and shoots were separated, they were washed under running water and fresh weight (g) was recorded. Dry weight (g) of shoot and root was measured too through drying the samples at 60°C in an electric oven (SANYO Drying Oven, MOV202, Japan) for a week.

### Assessment of Reproductive Traits and Fruit Yield

Flowering initiated on 19^th^ January 2023 and various reproductive data namely number of days required from transplanting to the first flowering, number of flower clusters plant^-1^, number of flowers and fruits cluster^-1^were noted against each replication under treatment. Harvesting was initiated on 23^rd^ February 2023 and continued till 9^th^ April 2023. Immediately after harvest, fruits were weighed (g) and averaged to get total fruit yield plant^-1^ by multiplying the individual fruit weight with the number of fruits cluster^-1^ and number of flower clusters plant^-1^.

### Statistical Analysis

Two-way analysis of variance (ANOVA) was performed for hypothesis test. Data were presented as the average of three replicates plus standard error (SE) where three observations were done per replication (n=3). The treatment means were separated by Fisher’s protected least significant difference (LSD) test, using a p-value of ≤ 0.05 to be statistically significant. In addition, correlation matrix and cluster analysis were conducted to examine the interrelationships among the studied dependent variables with respect to the fertilization and PGRs treatments where the strength of correlations among the properties were assessed according to Pearson’s correlation co-efficient. Afterwards, multivariate analysis like principal component analysis (PCA) was done to see more insight into the data matrix and sort out the most effective variables contribute in the total variations. Correlation matrix, cluster analysis and PCA were performed using different packages (agricole, facatominer, factoextra, ggplot2, corrplot) of ‘R’ program (version 4.1.2).

### Ethical statement

Tomato var. *BARI Tomato-14* (registered tomato variety of Bangladesh released by Bangladesh Agricultural Research Institute) was used as plant material and fertilizers at varied amounts and plant growth regulators namely GA_3_, NAA, 4-CPA and SA were used as treatment materials in the present study. Plastic pots were filled with soil media where tomato seedlings were transplanted and grown to obtain yield. The lab and field experiments in this study were carried out following guidelines and recommendations of “Biosafety Guidelines of Bangladesh” published by Ministry of Environment and Forest, Government of the People’s Republic of Bangladesh (2005).

## Results

### Weather Condition in Late Winter

Remarkable fluctuations and differences between maximum and minimum temperature as well as maximum and minimum relative humidity were noticed during the late winter growing period of tomato (Figure 1A and 1B). Both high and low temperature indexes showed gradual decreased down from 30.0 °C and 16.5 °C, respectively in the 1^st^ December, 2022 to 19.0 °C and 10.0 °C in the mid-January 2023. Thereafter, a steady increase in temperature was noticed up to the end of the harvest at April 2023 up to a maximum of 38.5 °C. Daily maximum temperature was noted below 30 °C up to the first week of February and from the second week of February 2023 maximum daily temperature climbed up. It was also noticed that temperature differentiation between high and low pick never went below 12.0 °C. Temperature variations in almost throughout the vegetative growth stage and reproductive phase were more than 15.0 °C (early-January to mid-March) (Figure 1A). Again, relative humidity report exhibited that except few deviations maximum relative humidity was more or less stable in 85-95 % index. While, extreme fluctuation was recorded in minimum daily relative humidity where the lowest humidity was observed 37 % in March 12, 2023. Throughout the growing period minimum relative humidity remained close to 60 % where relative minimum atmospheric humidity was recorded below 50 % during February and March 2023 (Figure 1B).

**Fig. 1.**
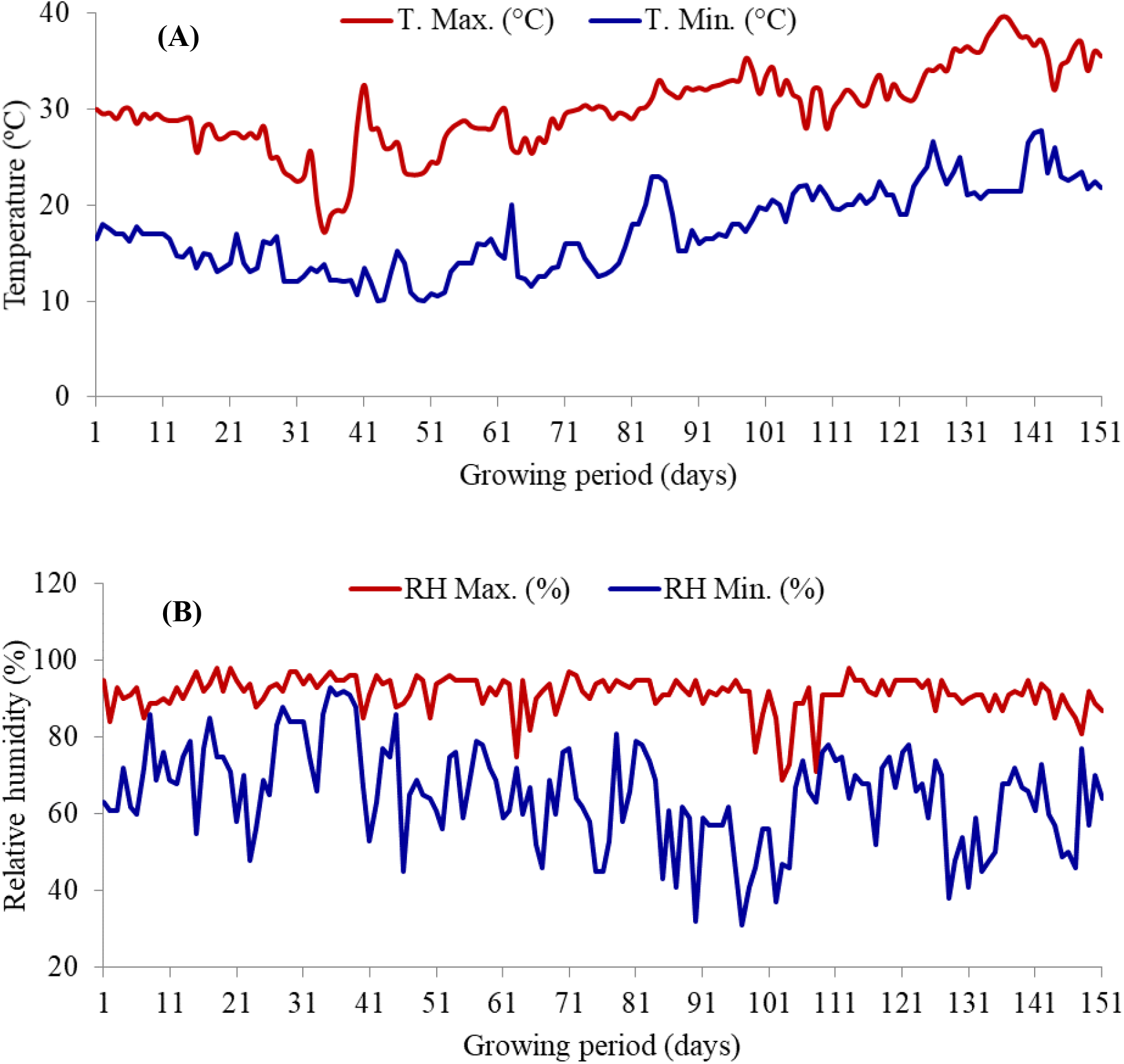
Temperature (A) and relative humidity (B) status of the experimental site during the growing period of tomato (December 01, 2022 to April 30, 2023). Here, T. Max. and T. Min. indicate maximum and minimum temperature, respectively and RH Max. and RH Min. represent maximum and minimum relative humidity, respectively.

### Plant Height

Plant height of tomato at 45 and 75 DAT was significantly (*p≤0.05*) influenced by the application of nutrients and plant growth regulators however plant height was not varied significantly at 20 DAT (Figure 2). Control plants (N_1_) for fertilizer treatment had shorter plants at both 45 and 75 DAT (38.45 cm and 63.41 cm, respectively) while the other tomato plants treated with 80 % to 110 % of FRG’2018 recommended fertilizer doses (N_2_ to N_5_) exhibited statistically similar and higher plant height compared to control (Figure 2A). Among the PGRs, maximum plant height was measured in P_2_ treatment at both 45 and 75 DAT (49.08 cm and 87.90 cm, respectively) which had statistical harmony with P_5_ (49.01 cm and 86.61 cm, respectively) followed by P_1_ treatment. Plant height at both the phases was recorded minimum in P_4_ treatment (31.35 cm and 46.78 cm, respectively) having statistical consistency with P_3_ treatment (33.27 cm and 47.67 cm, respectively) (Figure 2B). Further, interaction revealed that the tallest tomato plant was recorded in N_5_P_5_ at 45 DAT (50.70 cm) being statistically at par with N_1_P_5_, N_2_P_2_, N_2_P_5_, N_3_P_2_, N_3_P_5_, N_4_P_2_, N_4_P_5_ and N_5_P_2_ combinations and in N_3_P_2_ treatment at 75 DAT (90.63 cm) that was statistical parity with the P_5_ of N_2_, N_3_, N_4_ and N_5_ and P_2_ combination of N_2_, N_4_ and N_5_. Contrarily, the shortest plant at 45 DAT was noted in N_1_P_4_ (29.97 cm) having statistical unity with the interaction of P_3_ with N_1_, N_3_, N_4_ and N_5_ and P_4_ of N_2_, N_3_, N_4_ and N_5_. Again, at 75 DAT, interaction of P_3_ and P_4_ with N_1_, N_2_, N_3_, N_4_ and N_5_ had statistically similar plant height of which minimum been recorded in N_3_P_4_ combinations (46.13 cm). Plant height at 20 DAT didn’t vary significantly in interaction effect of fertilizers and PGRs (Table 1).

**Fig. 2.**
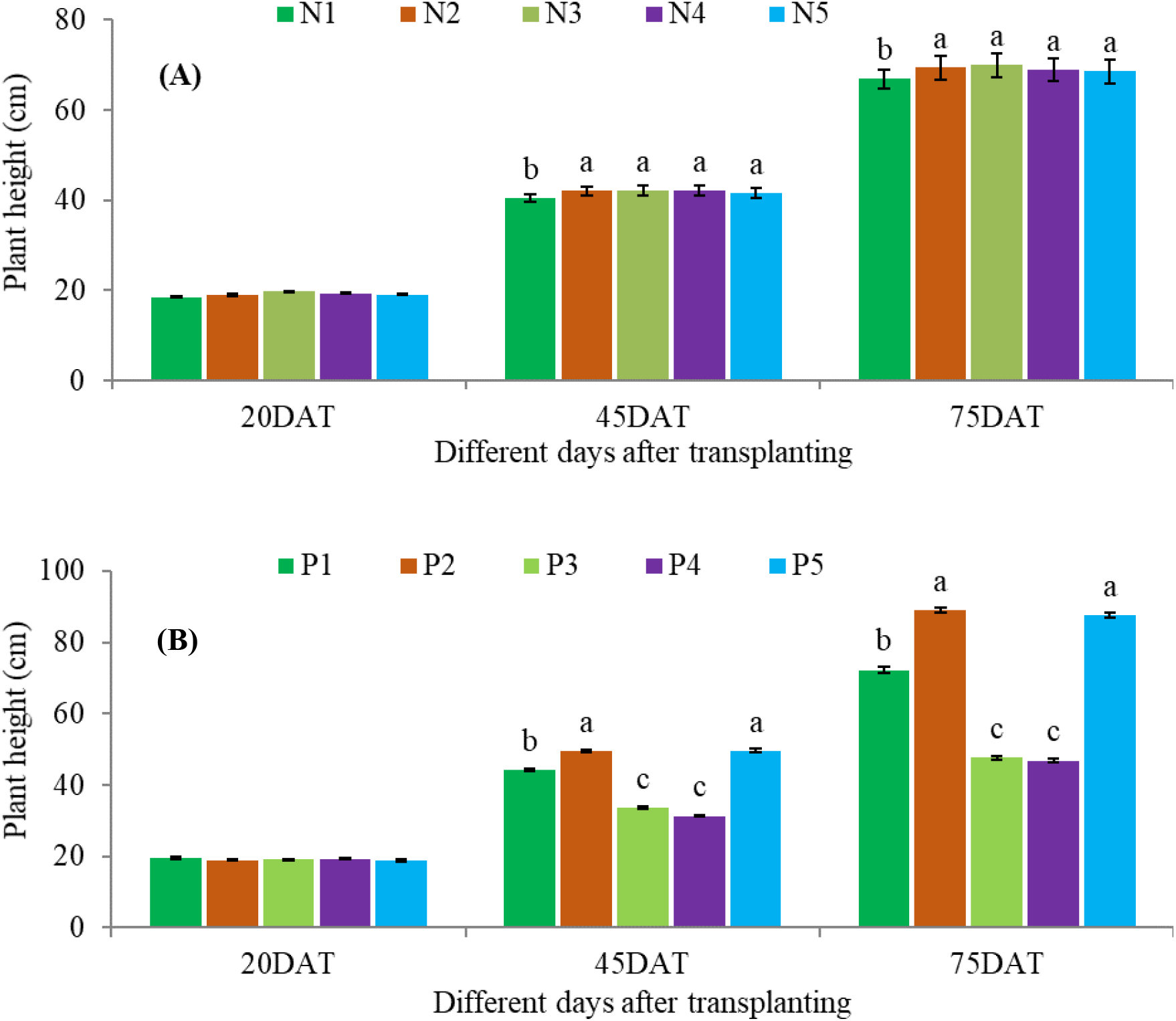
Plant height of tomato at different days after transplanting as influenced by the application of fertilizers (A) and plant growth regulators (B). Vertical bars on the top of the columns represent the standard errors of means of three replicates. Different letters indicate the statistical differences among the treatments at *p≤0.05*. Here, N_1_, N_2_, N_3_, N_4_ and N_5_ represent control (no fertilizer), 100, 110, 90 and 80 % of FRG’2018, respectively and P_1_, P_2_, P_3_, P_4_ and P_5_ indicate control (no PGR), GA_3_, NAA, 4-CPA and SA at 50 ppm, respectively.

**Table 1.** Interaction effect of fertilizer and PGR on plant height, base diameter and branch number of tomato at different days after transplanting (DAT)

| Treatment combination |  | Plant height at different DAT |  |  | Base diameter at different DAT |  |  | Branch number at different DAT |  |  |
| --- | --- | --- | --- | --- | --- | --- | --- | --- | --- | --- |
|  |  | 20 | 45 | 75 | 20 | 45 | 75 | 20 | 45 | 75 |
| N<sub>1</sub> | P<sub>1</sub> | 19.60 ± 1.39 | 39.10 ± 1.55de | 61.97 ± 2.48e | 0.57 ± 0.03 | 1.47 ± 0.12d-f | 2.13 ± 0.05fg | 1.30 ± 0.00 | 3.00 ± 0.17c-g | 4.00 ± 0.17ef |
|  | P<sub>2</sub> | 18.53 ± 1.36 | 45.73 ± 1.86bc | 80.97 ± 3.06b-d | 0.57 ± 0.03 | 1.73 ± 0.09a-c | 2.43 ± 0.01d-f | 1.43 ± 0.13 | 3.87 ± 0.30a-c | 5.23 ± 0.29a-d |
|  | P<sub>3</sub> | 18.07 ± 1.45 | 31.57 ± 1.07fg | 47.90 ± 3.85f | 0.57 ± 0.03 | 1.33 ± 0.09ef | 1.90 ± 0.06g | 1.30 ± 0.00 | 2.57 ± 0.30g | 3.67 ± 0.20ef |
|  | P<sub>4</sub> | 18.23 ± 0.93 | 29.97 ± 0.95g | 46.57 ± 3.45f | 0.57 ± 0.03 | 1.30 ± 0.06ef | 1.83 ± 0.09g | 1.57 ± 0.13 | 2.33 ± 0.20g | 3.67 ± 0.38ef |
|  | P<sub>5</sub> | 18.40 ± 1.47 | 45.87 ± 1.87a-c | 79.67 ± 2.95cd | 0.57 ± 0.03 | 1.70 ± 0.12a-d | 2.37 ± 0.15ef | 1.57 ± 0.13 | 3.53 ± 0.39a-f | 5.13 ± 0.30a-d |
| N<sub>2</sub> | P<sub>1</sub> | 18.47 ± 1.84 | 43.80 ± 1.94cd | 72.00 ± 3.53d | 0.57 ± 0.03 | 1.53 ± 0.09b-e | 2.43 ± 0.09d-f | 1.43 ± 0.13 | 3.57 ± 0.70a-f | 4.57 ± 0.47c-e |
|  | P<sub>2</sub> | 18.40 ± 1.24 | 50.13 ± 1.64ab | 89.93 ± 3.58ab | 0.57 ± 0.03 | 1.77 ± 0.09ab | 2.77 ± 0.02a-c | 1.43 ± 0.13 | 3.67 ± 0.20a-e | 5.87 ± 0.30ab |
|  | P<sub>3</sub> | 19.47 ± 0.65 | 35.17 ± 1.23ef | 48.43 ± 2.69f | 0.57 ± 0.03 | 1.30 ± 0.06ef | 2.00 ± 0.12g | 1.33 ± 0.03 | 2.57 ± 0.30g | 3.47 ± 0.39f |
|  | P<sub>4</sub> | 19.53 ± 0.71 | 33.03 ± 1.26fg | 47.33 ± 3.44f | 0.57 ± 0.03 | 1.33 ± 0.09ef | 1.97 ± 0.09g | 1.30 ± 0.00 | 2.77 ± 0.23e-g | 3.43 ± 0.43f |
|  | P<sub>5</sub> | 18.97 ± 1.74 | 48.17 ± 2.73a-c | 89.20 ± 3.52ab | 0.60 ± 0.06 | 1.77 ± 0.09ab | 2.70 ± 0.12a-d | 1.57 ± 0.13 | 4.00 ± 0.40ab | 5.67 ± 0.49ab |
| N<sub>3</sub> | P<sub>1</sub> | 20.67 ± 0.73 | 45.83 ± 2.00bc | 76.83 ± 3.09d | 0.57 ± 0.03 | 1.60 ± 0.06a-d | 2.50 ± 0.01b-e | 1.57 ± 0.01 | 4.00 ± 0.17ab | 5.67 ± 0.20ab |
|  | P<sub>2</sub> | 20.10 ± 0.87 | 50.33 ± 2.07ab | 90.63 ± 2.95a | 0.57 ± 0.03 | 1.77 ± 0.09ab | 2.80 ± 0.12ab | 1.57 ± 0.13 | 4.00 ± 0.40ab | 6.13 ± 0.43a |
|  | P<sub>3</sub> | 19.30 ± 1.10 | 34.10 ± 1.49fg | 47.63 ± 3.06f | 0.53 ± 0.03 | 1.27 ± 0.09f | 1.97 ± 0.09g | 1.30 ± 0.00 | 2.87 ± 0.30d-g | 3.67 ± 0.33ef |
|  | P<sub>4</sub> | 19.10 ± 0.60 | 31.00 ± 1.59fg | 46.13 ± 3.23f | 0.57 ± 0.03 | 1.27 ± 0.09f | 1.90 ± 0.06g | 1.43 ± 0.13 | 2.67 ± 0.20fg | 3.57 ± 0.13ef |
|  | P<sub>5</sub> | 19.60 ± 0.78 | 49.97 ± 2.08ab | 88.77 ± 2.87a-c | 0.57 ± 0.03 | 1.77 ± 0.12ab | 2.77 ± 0.15a-c | 1.43 ± 0.13 | 3.87 ± 0.30a-c | 5.47 ± 0.23a-c |
| N<sub>4</sub> | P<sub>1</sub> | 19.10 ± 0.59 | 45.73 ± 1.41bc | 73.23 ± 4.04d | 0.57 ± 0.03 | 1.50 ± 0.06c-f | 2.47 ± 0.09c-e | 1.43 ± 0.13 | 3.23 ± 0.23b-g | 4.57 ± 0.30c-e |
|  | P<sub>2</sub> | 20.37 ± 0.96 | 49.67 ± 2.41ab | 89.20 ± 3.44ab | 0.57 ± 0.03 | 1.80 ± 0.12a | 2.77 ± 0.15a-c | 1.40 ± 0.10 | 3.77 ± 0.23a-d | 5.90 ± 0.20ab |
|  | P<sub>3</sub> | 19.53 ± 0.61 | 32.83 ± 1.60fg | 47.30 ± 2.70f | 0.60 ± 0.00 | 1.27 ± 0.09f | 1.93 ± 0.15g | 1.43 ± 0.13 | 3.00 ± 0.17c-g | 3.87 ± 0.30ef |
|  | P<sub>4</sub> | 20.07 ± 1.11 | 31.93 ± 1.31fg | 47.60 ± 2.80f | 0.57 ± 0.03 | 1.27 ± 0.09f | 1.93 ± 0.12g | 1.33 ± 0.03 | 3.00 ± 0.17c-g | 4.00 ± 0.40ef |
|  | P<sub>5</sub> | 17.90 ± 0.92 | 50.33 ± 2.59ab | 86.83 ± 3.56a-c | 0.57 ± 0.03 | 1.77 ± 0.09ab | 2.77 ± 0.09a-c | 1.43 ± 0.13 | 3.67 ± 0.33a-e | 5.10 ± 0.10b-d |
| N<sub>5</sub> | P<sub>1</sub> | 19.63 ± 0.75 | 44.17 ± 1.33c | 71.97 ± 3.85d | 0.57 ± 0.03 | 1.50 ± 0.06c-f | 2.43 ± 0.09d-f | 1.43 ± 0.13 | 3.20 ± 0.49b-g | 4.43 ± 0.59d-f |
|  | P<sub>2</sub> | 17.53 ± 0.58 | 49.53 ± 1.85ab | 88.77 ± 3.12a-c | 0.57 ± 0.03 | 1.77 ± 0.09ab | 2.83 ± 0.09a | 1.43 ± 0.13 | 4.23 ± 0.23a | 6.10 ± 0.49ab |
|  | P<sub>3</sub> | 19.07 ± 0.83 | 32.70 ± 1.39fg | 47.10 ± 2.51f | 0.57 ± 0.03 | 1.30 ± 0.06ef | 1.93 ± 0.09g | 1.30 ± 0.00 | 2.77 ± 0.23e-g | 3.67 ± 0.20ef |
|  | P<sub>4</sub> | 19.60 ± 1.25 | 30.80 ± 0.76fg | 46.27 ± 2.95f | 0.53 ± 0.03 | 1.33 ± 0.09ef | 1.93 ± 0.09g | 1.43 ± 0.13 | 2.77 ± 0.23e-g | 3.67 ± 0.33ef |
|  | P<sub>5</sub> | 19.60 ± 0.78 | 50.70 ± 1.77a | 88.57 ± 3.08a-c | 0.57 ± 0.03 | 1.73 ± 0.12a-c | 2.77 ± 0.12a-c | 1.37 ± 0.07 | 3.90 ± 0.49a-c | 5.43 ± 0.59a-d |
| LS |  | ns | * | * | ns | * | * | ns | * | * |

Values are means ± standard errors of three replications. Different letters within the column indicate statistically significant differences among the treatments according to LSD at $p \leq 0.05$ . Here, N<sub>1</sub>, N<sub>2</sub>, N<sub>3</sub>, N<sub>4</sub> and N<sub>5</sub> represent control (no fertilizer), 100, 110, 90 and 80 % of FRG'2018, respectively and P<sub>1</sub>, P<sub>2</sub>, P<sub>3</sub>, P<sub>4</sub> and P<sub>5</sub> indicate control (no PGR), GA<sub>3</sub>, NAA, 4-CPA and SA at 50 ppm, respectively. LS: Level of significance, ns: Not significant, \*: Significant at 5 % level of probability

### Base diameter

Main effect of fertilizer exhibited non-significant variation in base diameter of tomato at 20 and 45 DAT. But base diameter at 75 DAT varied statistically where maximum base diameter was noted in N_3_ (2.39 cm) being statistically similar with N_2_ (2.37 cm), N_4_ (2.37 cm) and N_5_ (2.38 cm) while minimum base diameter was measured in control (2.13 cm) (Figure 3A). Again, base diameter differed significantly among the PGR treatments with the exception at 20 DAT (Figure 3B). At 45 and 75 DAT, maximum base diameter was recorded in P_2_ PGR (1.77 cm and 2.72 cm, respectively) which had statistical uniformity with P_5_ (1.75 cm and 2.67 cm, respectively) while minimum base diameter was measured in P_3_ at 45 DAT (1.29 cm) and in P_5_ at 75 DAT (1.91 cm). Interaction effect also showed significant difference in base diameter at 45 and 75 DAT (Table 1). At 45 DAT, N_3_P_3_, N_3_P_4_, N_4_P_3_ and N_4_P_4_ treatment combinations had significantly minimum same base diameter (1.27 cm) which had statistical similarity with N_1_, N_2_ and N_5_ interacted P_3_ and P_4_ as well as N_1_, N_4_ and N_5_ interacted P_1_. Whereas, maximum base diameter at 45 DAT was noted in N_4_P_2_ (1.80 cm) having statistical conformity with nutrient and P_2_ and P_5_ interaction. Again, at 75 DAT, significantly maximum base diameter was recorded in N_5_P_2_ (2.83 cm) being identical to P_2_ and P_5_ interaction of N_2_, N_3_, N_4_ and N_5_ treatments while minimum base diameter in N_1_P_4_ (1.83 cm) and P_3_ and P_4_ interaction of nutrient doses had statistical parity with that treatment combination (Table 1).

**Fig. 3.**
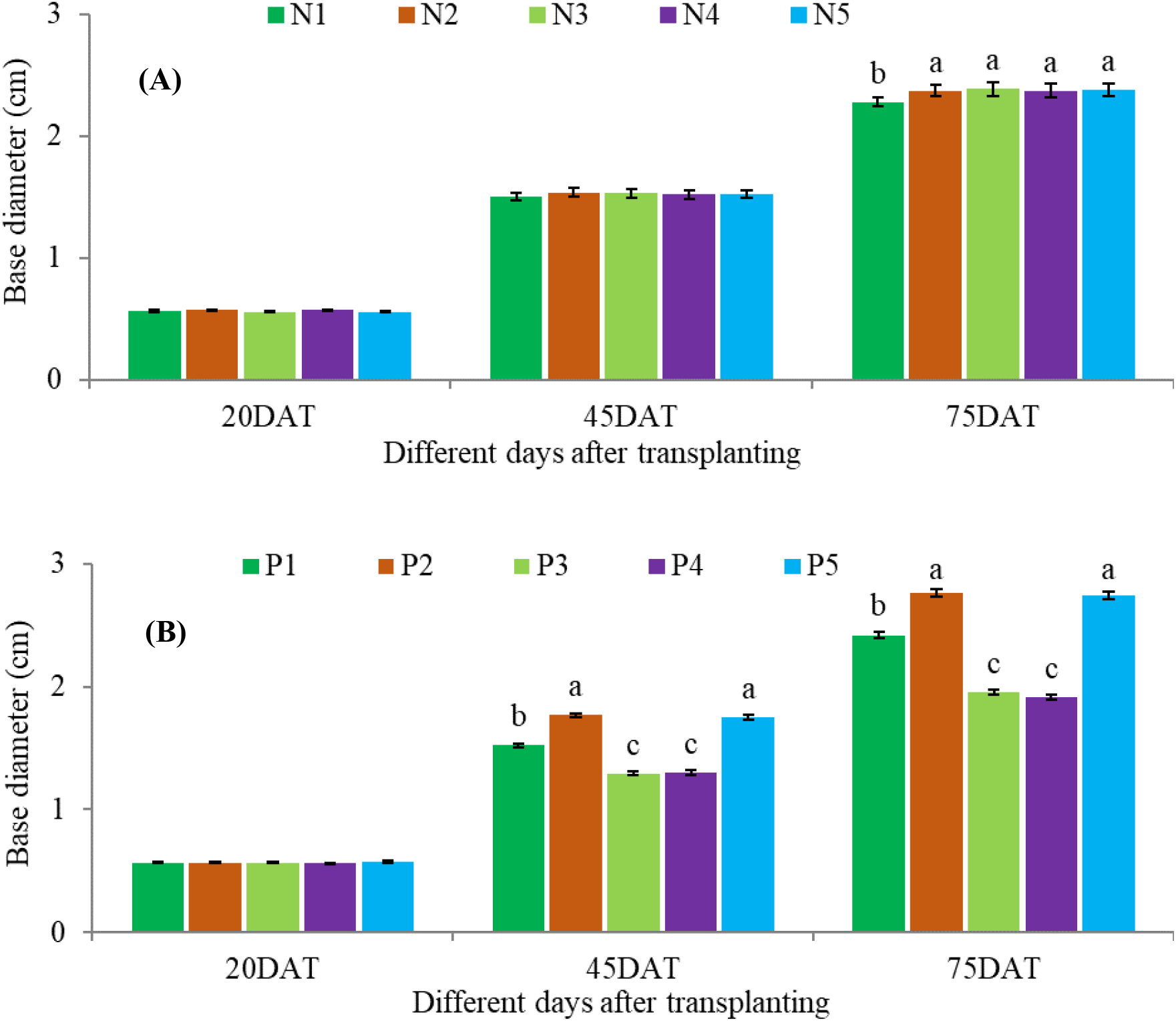
Base diameter of tomato at different days after transplanting as influenced by the application of fertilizers (A) and plant growth regulators (B). Vertical bars on the top of the columns represent the standard errors of means of three replicates. Different letters indicate the statistical differences among the treatments at *p≤0.05*. Here, N_1_, N_2_, N_3_, N_4_ and N_5_ represent control (no fertilizer), 100, 110, 90 and 80 % of FRG’2018, respectively and P_1_, P_2_, P_3_, P_4_ and P_5_ indicate control (no PGR), GA_3_, NAA, 4-CPA and SA at 50 ppm, respectively.

### Number of Branches

Number of branches didn’t vary with the variation in fertilizer doses rather it was varied remarkably due to PGR variations (Figure 4). At 45 DAT, maximum number of branches was found in P_3_ treatment (3.91 plant^-1^) having statistical conformity with P_5_ while minimum number of branches was counted in P_4_ (2.71 plant^-1^) and P_3_ (2.75 plant^-1^). Statistical superiority in branch number at 75 DAT was observed again in P_3_ treatment (5.85 plant^-1^) whereas P_3_ and P_4_ had statistically same and inferior branch number (3.67 plant^-1^) (Figure 4B). It was observed from the interaction effect that N_1_, N_2_, N_3_, N_4_ and N_5_ interacted P_2_ and P_5_ along with N_3_P_1_ expressed statistical similarity for number of branches plant-1 where N_5_P_2_ and N_3_P_2_ had maximum number of branches (4.23 and 6.13 branches plant^-1^, respectively) at 45 and 75 DAT. The rest P_1_, P_3_ and P_4_ interaction of all the nutrient treatment had statistically uniform and minimum number of branches plant^-1^ (Table 1).

**Fig. 4.**
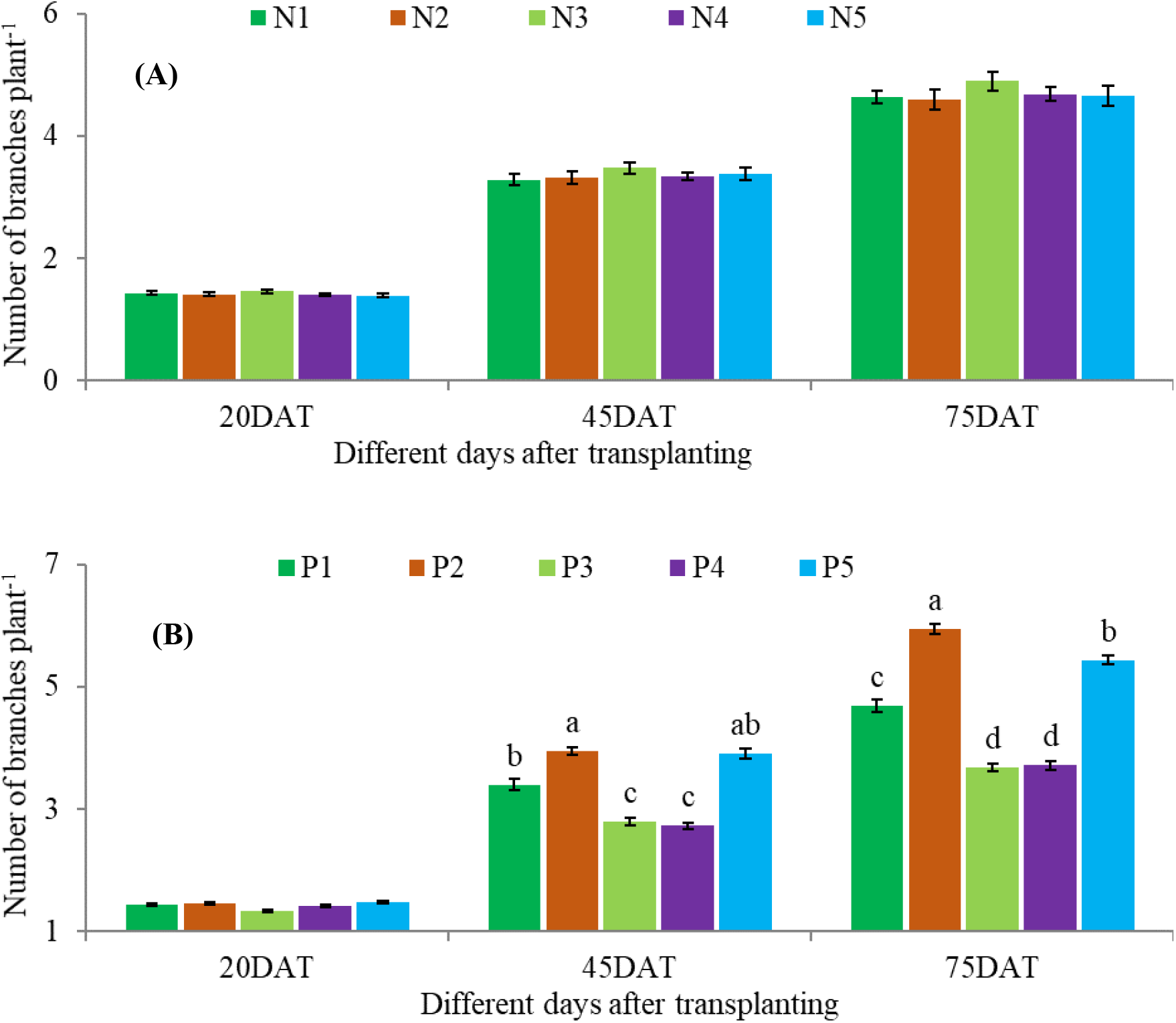
Number of branches plant^-1^ of tomato at different days after transplanting as influenced by the application of fertilizers (A) and plant growth regulators (B). Vertical bars on the top of the columns represent the standard errors of means of three replicates. Different letters indicate the statistical differences among the treatments at *p≤0.05*. Here, N_1_, N_2_, N_3_, N_4_ and N_5_ represent control (no fertilizer), 100, 110, 90 and 80 % of FRG’2018, respectively and P_1_, P_2_, P_3_, P_4_ and P_5_ indicate control (no PGR), GA_3_, NAA, 4-CPA and SA at 50 ppm, respectively.

### Number of Leaves

Number of leaves plant^-1^ of tomato at 45 and 75 DAT was significantly (*p≤0.05*) influenced by the application of nutrients and PGRs but non-significant variation was evident at 20 DAT (Figure 5). Among the fertilizer doses, maximum number of leaves at 45 and 75 DAT was counted in N_3_ treatment (22.36 and 36.44 plant^-1^) having statistical consistency with N_2_ and N_4_ treatments while minimum number of leaves at those two stages was noted in N_1_ fertilizer dose (19.78 and 31.44 plant^-1^) (Figure 5A). Again, P_2_ among the studied PGR exhibited statistically superior number of leaves at both 45 and 75 DAT (27.69 and 47.13 plant^-1^) being different from all other treatments while statistically inferior number of leaves at both 45 and 75 DAT was recorded in P_4_ treatment (16.44 and 25.77 plant^-1^) being statistically uniform with P_3_ at 45 DAT and with P_1_ and P_3_ at 75 DAT (Figure 5B). Moreover, interaction revealed that except N_3_P_1_, PGRs like P_1_, P_3_ and P_4_ interacted with N_1_ to N_5_ nutrient doses had statistical parity for number of leaves plant^-1^ at 45 DAT. Of them, plants under N_5_P_4_ interaction produced the lowest number of leaves (16.00 plant^-1^). On the contrary, N_2_P_2_ combination had the highest number of leaves (29.00 plant^-1^) at 75 DAT and P_2_ and P_3_ combined with N_3_ and N_4_ and P_1_ interacted with N_1_, N_2_ and N_5_ had statistical consistency among them. Reversely, N_1_P_4_ treated plants obtained the lowest number of leaves (21.00 plant^-1^) at 75 DAT having statistical uniformity with N_1_P_1_, N_1_P_3_, N_2_P_3_, N_4_P_3_, N_4_P_4_, N_5_P_1_, N_5_P_3_ and N_5_P_4_ combinations (Table 2).

**Fig. 5.**
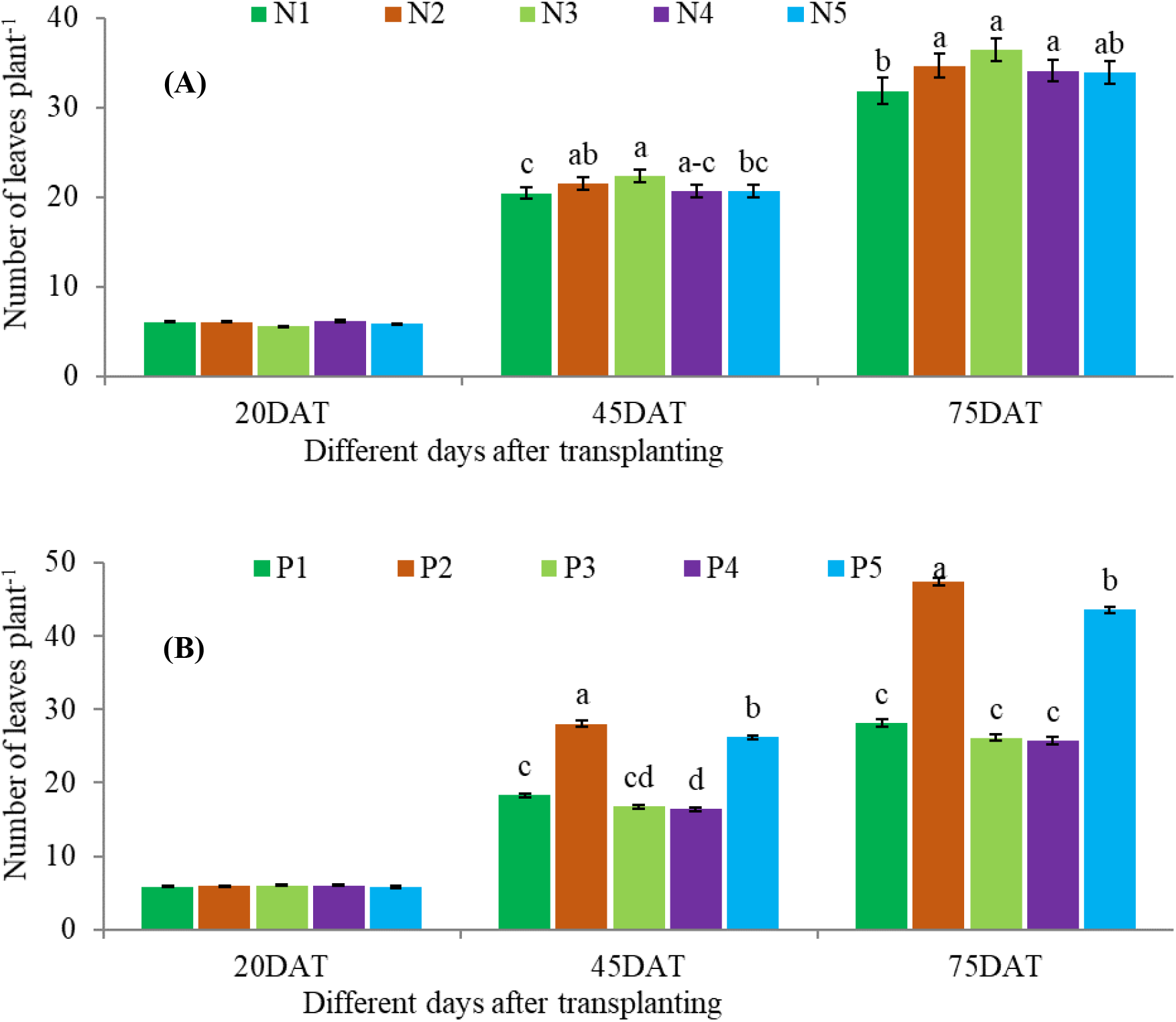
Number of leaves plant^-1^ of tomato at different days after transplanting as influenced by the application of fertilizers (A) and plant growth regulators (B). Vertical bars on the top of the columns represent the standard errors of means of three replicates. Different letters indicate the statistical differences among the treatments at *p≤0.05*. Here, N_1_, N_2_, N_3_, N_4_ and N_5_ represent control (no fertilizer), 100, 110, 90 and 80 % of FRG’2018, respectively and P_1_, P_2_, P_3_, P_4_ and P_5_ indicate control (no PGR), GA_3_, NAA, 4-CPA and SA at 50 ppm, respectively.

**Table 2.**
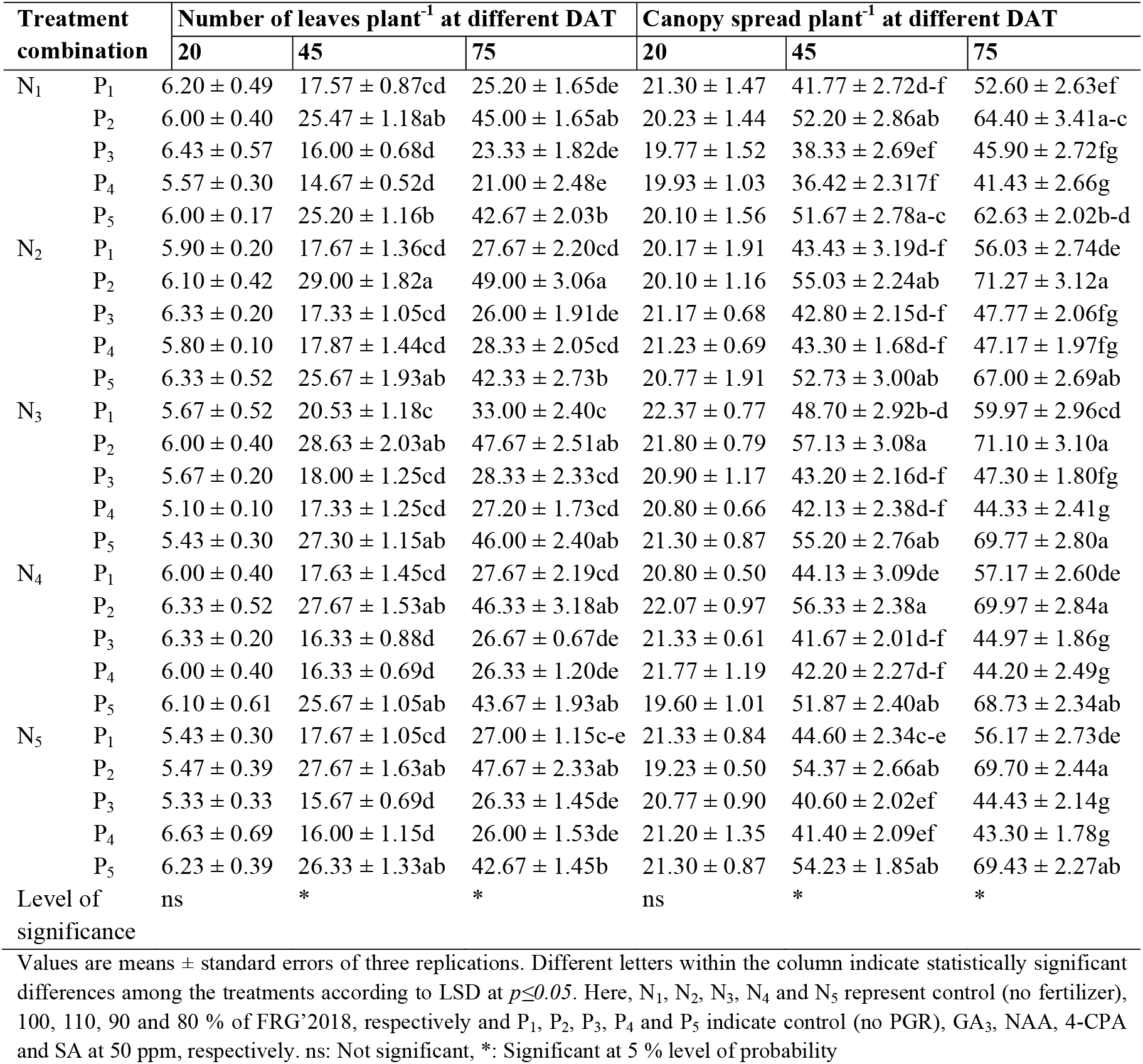
Fertilizer and PGR interactions influencing the number of leaves and canopy spread plant^-1^ of tomato at different days after transplanting (DAT)

### Canopy Spread

Fertilizer doses and PGRs implied no statistical variation on canopy spread at 20 DAT considering their main and interaction effect, but significant (*p≤0.05*) differences were noticed at 45 and 75 DAT. Tomato plants treated with N_3_ fertilizer dose attained maximum canopy spread at both 45 and 75 DAT (49.27 and 58.49 cm plant^-1^, respectively). Rest of the nutrient doses except control (N_1_) got statistical harmony with N_3_ treatment. Minimum canopy spread was estimated in N_1_ fertilizer treated plants (44.09 cm and 53.39 cm at 45 and 75 DAT, respectively) (Figure 6A). On the other side, P_2_ PGR resulted in maximum canopy spread of tomato at 45 and 75 DAT (55.01 and 69.29 cm, respectively) which had statistical uniformity with P_5_ treatment (53.14 and 67.51 cm, respectively) while canopy spread measured the lowest in P_4_ (41.10 and 44.09 cm, respectively) having statistical parity with P3 treatment (41.32 and 46.07 cm, respectively) (Figure 6B). Further, among nutrient-PGR interactions, canopy spread of tomato was estimated the utmost in N_3_P_2_ combinations at 45 DAT (57.13 cm) and in N_2_P_2_ at 75 DAT (71.27 cm). Interactions like N_1_P_2_, N_1_P_5_, N_2_P_2_, N_2_P_5_, N_3_P_5_, N_4_P_2_, N_4_P_5_, N_5_P_2_ and N_5_P_5_ at 45 DAT and N_1_P_2_, N_2_P_5_, N_3_P_5_, N_4_P_2_, N_4_P_5_, N_5_P_2_ and N_5_P_5_ at 75 DAT had statistical unity with the best combination. On the contrary, plants under N_1_P_4_ combination attained minimum canopy spread at both 45 and 75 DAT (36.42 and 41.43 cm, respectively) which got statistical similarity with N_1_P_1_, N_1_P_3_, N_2_P_1_, N_2_P_3_, N_2_P_4_, N_3_P_3_, N_3_P_4_, N_4_P_3_, N_4_P_4_, N_5_P_3_ and N_5_P_4_ combinations at 45 DAT and with N_1_P_3_, N_2_P_3_, N_2_P_4_, N_3_P_3_, N_3_P_4_, N_4_P_3_, N_4_P_4_, N_5_P_3_ and N_5_P_4_ combinations at 75 DAT (Table 2).

**Fig. 6.**
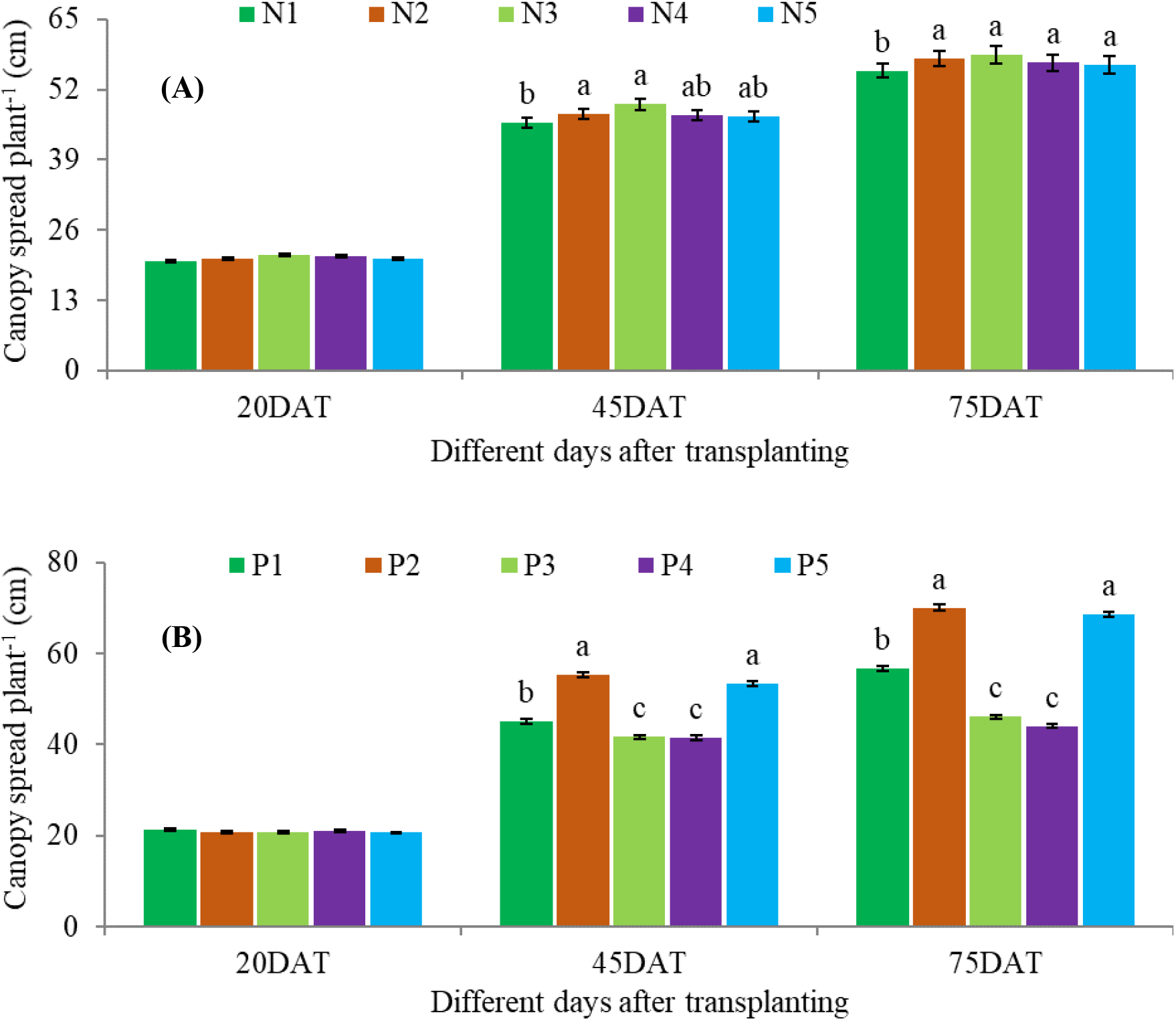
Canopy spread plant^-1^ of tomato at different days after transplanting as influenced by the application of fertilizers (A) and plant growth regulators (B). Vertical bars on the top of the columns represent the standard errors of means of three replicates. Different letters indicate the statistical differences among the treatments at *p≤0.05*. Here, N_1_, N_2_, N_3_, N_4_ and N_5_ represent control (no fertilizer), 100, 110, 90 and 80 % of FRG’2018, respectively and P_1_, P_2_, P_3_, P_4_ and P_5_ indicate control (no PGR), GA_3_, NAA, 4-CPA and SA at 50 ppm, respectively.

### Internode Length

Internode length of tomato varied non-significantly with fertilizer application at different doses but variation was significant in terms of PGR treatment (Table 3). Due to fertilizer feeding internode length of tomato ranged between 4.92 cm and 5.13 cm. In case of PGR treatment, long (6.17 cm) internode was recorded in P_2_ followed by P_5_ treatment (5.53 cm). While, short internode was measured in P_4_ treatment (4.20 cm) being statistically identical with P_3_ treatment (4.32 cm). Interaction of nutrient and PGR also revealed significant variation in internode length of tomato where plants under N_2_P_2_ treatment combination had the longest internode (6.53 cm) having statistical parity with N_1_P_2_, N_3_P_2_, N_3_P_5_, N_4_P_2_, N_5_P_2_ and N_5_P_5_ combinations. Conversely, the shortest internode was noticed in N_1_P_4_ interaction (4.03 cm) which had statistical similarity with N_1_P_1_, N_1_P_3_, N_2_P_3_, N_2_P_4_, N_3_P_3_, N_3_P_4_, N_4_P_3_, N_4_P_4_, N_5_P_3_ and N_5_P_4_ treatments (Table 4).

**Table 3.** Number of leaflets leaf-1, internode length, leaf area, leaf SPAD value and fresh and dry weight of shoot and root of tomato as influenced by the application of fertilizers and plant growth regulators.

| Treatment | No. of leaflets<br>leaf<sup>1</sup> | Internode<br>length (cm) | Leaf area (cm<sup>2</sup>) | Leaf SPAD<br>value | Shoot weight (g) |  | Root weight (g) |  |
| --- | --- | --- | --- | --- | --- | --- | --- | --- |
|  |  |  |  |  | Fresh | Dry | Fresh | Dry |
| Fertilizer dose |  |  |  |  |  |  |  |  |
| N<sub>1</sub> | 7.94 ± 0.37b | 4.92 ± 0.22 | 286.33 ± 8.10b | 48.59 ± 1.21 | 265.61 ± 11.6b | 62.34 ± 2.20b | 33.84 ± 1.64b | 18.87 ± 0.50 |
| N<sub>2</sub> | 8.05 ± 0.40b | 5.08 ± 0.28 | 305.54 ± 7.07a | 50.44 ± 1.14 | 282.37 ± 10.9a | 66.17 ± 1.78a | 35.61 ± 1.46a | 19.33 ± 0.42 |
| N<sub>3</sub> | 8.31 ± 0.46ab | 5.13 ± 0.22 | 306.13 ± 13.9a | 51.34 ± 1.09 | 283.94 ± 13.8a | 67.91 ± 2.23a | 35.81 ± 1.74a | 19.16 ± 0.54 |
| N<sub>4</sub> | 8.47 ± 0.37a | 5.08 ± 0.22 | 300.33 ± 12a | 49.94 ± 0.95 | 278.38 ± 13.9a | 66.54 ± 1.91a | 35.30 ± 1.76a | 19.46 ± 0.40 |
| N<sub>5</sub> | 8.58 ± 0.46a | 5.06 ± 0.22 | 300.16 ± 12.5a | 50.21 ± 0.96 | 277.60 ± 14.5a | 65.92 ± 1.98a | 35.25 ± 1.77a | 19.00 ± 0.48 |
| LS | * | ns | * | ns | ** | * | * | ns |
| Plant growth regulator |  |  |  |  |  |  |  |  |
| P<sub>1</sub> | 8.51 ± 0.15c | 5.06 ± 0.10c | 314.41 ± 5.99b | 49.42 ± 1.02bc | 289.98 ± 4.78b | 66.59 ± 1.34b | 37.19 ± 0.57b | 19.21 ± 0.36b |
| P<sub>2</sub> | 10.19 ± 0.13a | 6.17 ± 0.14a | 337.66 ± 5.24a | 53.08 ± 1.00a | 328.13 ± 4.46a | 73.20 ± 1.28a | 41.47 ± 0.48a | 20.77 ± 0.26a |
| P<sub>3</sub> | 6.38 ± 0.20e | 4.32 ± 0.13d | 270.77 ± 7.18c | 47.87 ± 0.83c | 229.52 ± 4.61c | 58.74 ± 0.92c | 28.65 ± 0.50c | 17.79 ± 0.29c |
| P<sub>4</sub> | 6.91 ± 0.15d | 4.20 ± 0.13d | 247.89 ± 6.47d | 48.15 ± 0.83c | 216.82 ± 3.55d | 57.86 ± 0.95c | 27.39 ± 0.42c | 17.43 ± 0.28c |
| P<sub>5</sub> | 9.36 ± 0.16b | 5.53 ± 0.12b | 327.75 ± 6.64ab | 52.01 ± 1.03ab | 323.45 ± 3.82a | 72.49 ± 1.15a | 41.10 ± 0.52a | 20.61 ± 0.29a |
| LS | ** | ** | ** | ** | ** | ** | ** | ** |
| CV (%) | 6.79 | 10.72 | 6.22 | 7.49 | 4.88 | 6.44 | 5.01 | 6.11 |

Values are means ± standard errors of three replications. Different letters within the column indicate statistically significant differences among the treatments according to LSD at $p \leq 0.05$ . Here, N<sub>1</sub>, N<sub>2</sub>, N<sub>3</sub>, N<sub>4</sub> and N<sub>5</sub> represent control (no fertilizer), 100, 110, 90 and 80 % of FRG'2018, respectively and P<sub>1</sub>, P<sub>2</sub>, P<sub>3</sub>, P<sub>4</sub> and P<sub>5</sub> indicate control (no PGR), GA<sub>3</sub>, NAA, 4-CPA and SA at 50 ppm, respectively. LS: Level of significance, \* and \*\*: Significant at 5 % and 1 % level of probability, respectively, ns: Not significant

**Table 4.** Interactive influence of fertilizer and PGR on number of leaflets leaf-1, internode internode length, leaf area, leaf SPAD value and fresh and dry weight of shoot and root of tomato.

| Treatment combination |  | No. of leaflets leaf<sup>1</sup> | Internode length (cm) | Leaf area (cm<sup>2</sup>) | SPAD value | Shoot weight (g) |  | Root weight (g) |  |
| --- | --- | --- | --- | --- | --- | --- | --- | --- | --- |
|  |  |  |  |  |  | Fresh | Dry | Fresh | Dry |
| N<sub>1</sub> | P<sub>1</sub> | 8.50 ± 0.17fg | 4.87 ± 0.42f-j | 286.40 ± 11.58g-j | 49.50 ± 2.68 | 273.89 ± 14.32g | 64.27 ± 4.29e-g | 35.66 ± 1.59f | 19.32 ± 1.06b-g |
|  | P<sub>2</sub> | 9.67 ± 0.07b-d | 6.00 ± 0.38a-d | 321.11 ± 9.84c-f | 51.77 ± 2.67 | 305.34 ± 7.29d-f | 68.70 ± 3.61b-e | 39.69 ± 1.17b-e | 20.49 ± 0.60a-c |
|  | P<sub>3</sub> | 5.93 ± 0.29k | 4.40 ± 0.26h-j | 264.40 ± 10.66jk | 45.63 ± 2.83 | 220.10 ± 5.61i-k | 56.04 ± 3.04hi | 27.45 ± 1.44h-j | 17.75 ± 1.06g-j |
|  | P<sub>4</sub> | 6.93 ± 0.37ij | 4.03 ± 0.09j | 248.43 ± 5.18kl | 45.17 ± 2.18 | 214.52 ± 4.67jk | 53.47 ± 2.75i | 26.49 ± 0.85ij | 16.46 ± 0.10j |
|  | P<sub>5</sub> | 8.67 ± 0.24e-g | 5.30 ± 0.4c-g | 311.31 ± 6.89d-h | 50.90 ± 2.40 | 314.22 ± 6.42b-e | 69.24 ± 3.92a-e | 39.90 ± 1.56b-d | 20.32 ± 0.29a-d |
| N<sub>2</sub> | P<sub>1</sub> | 8.47 ± 0.29fg | 5.13 ± 0.13d-h | 310.94 ± 10.6d-h | 47.50 ± 3.95 | 285.94 ± 4.94fg | 64.67 ± 3.90d-g | 36.46 ± 0.76f | 18.83 ± 0.98c-h |
|  | P<sub>2</sub> | 9.80 ± 0.31a-c | 6.53 ± 0.41a | 326.65 ± 6.67b-f | 53.03 ± 2.13 | 327.29 ± 8.89a-d | 73.43 ± 1.89a-c | 41.36 ± 0.98a-c | 20.79 ± 0.27ab |
|  | P<sub>3</sub> | 6.13j ± 0.84k | 4.23 ± 0.38ij | 282.99 ± 11.2h-j | 49.33 ± 1.20 | 239.70 ± 5.50hi | 60.43 ± 2.05f-h | 29.64 ± 0.38gh | 18.42 ± 0.57d-i |
|  | P<sub>4</sub> | 6.93 ± 0.18ij | 4.03 ± 0.52j | 274.43 ± 6.23i-k | 49.73 ± 1.14 | 233.68 ± 3.07h-j | 59.93 ± 0.41f-i | 29.14 ± 0.54g-i | 17.79 ± 0.06g-j |
|  | P<sub>5</sub> | 8.93 ± 0.18c-f | 5.47 ± 0.29b-f | 332.68 ± 9.24a-e | 52.60 ± 3.38 | 325.24 ± 1.99a-d | 72.37 ± 2.19a-c | 41.46 ± 0.30a-c | 20.81 ± 0.91ab |
| N<sub>3</sub> | P<sub>1</sub> | 8.87 ± 0.47d-f | 5.30 ± 0.26c-g | 342.52 ± 9.79a-c | 52.33 ± 1.74 | 310.56 ± 7.28c-e | 70.67 ± 2.99a-e | 39.52 ± 1.12c-e | 19.80 ± 1.11a-f |
|  | P<sub>2</sub> | 10.40 ± 0.23ab | 6.07 ± 0.35a-c | 357.33 ± 13.97a | 54.93 ± 2.58 | 343.20 ± 8.21a | 76.17 ± 3.08a | 42.82 ± 0.81a | 21.23 ± 0.86ab |
|  | P<sub>3</sub> | 6.47 ± 0.29jk | 4.30 ± 0.26h-j | 306.13 ± 15.85e-h | 48.37 ± 2.80 | 249.97 ± 12.71h | 60.00 ± 1.99f-i | 30.76 ± 1.05g | 17.67 ± 0.46g-j |
|  | P<sub>4</sub> | 6.27 ± 0.29jk | 4.27 ± 0.19h-j | 225.03 ± 27.33l | 47.77 ± 0.73 | 204.88 ± 11.86k | 58.00 ± 0.96g-i | 26.12 ± 1.36j | 16.70 ± 0.27ij |
|  | P<sub>5</sub> | 9.53 ± 0.35b-e | 5.73 ± 0.23a-f | 299.61 ± 12.05f-i | 53.30 ± 2.00 | 311.11 ± 10.24c-e | 74.70 ± 3.02ab | 39.84 ± 1.31b-d | 20.37 ± 0.87a-c |
| N<sub>4</sub> | P<sub>1</sub> | 7.87 ± 0.24gh | 5.03 ± 0.18e-i | 318.56 ± 4.79c-f | 47.93 ± 1.21 | 292.15 ± 8.49e-g | 66.60 ± 1.30c-f | 37.38 ± 1.07d-f | 19.32 ± 0.57b-g |
|  | P<sub>2</sub> | 10.40 ± 0.12ab | 6.23 ± 0.23ab | 344.00 ± 12.25a-c | 55.03 ± 2.41 | 333.93 ± 7.8ab | 75.80 ± 3.12a | 41.93 ± 1.01a-c | 21.49 ± 0.19a |
|  | P<sub>3</sub> | 6.93 ± 0.12ij | 4.43 ± 0.43g-j | 251.63 ± 8.48kl | 48.30 ± 0.81 | 220.02 ± 7.85i-k | 59.20 ± 1.31g-i | 27.84 ± 0.80h-j | 18.17 ± 0.38e-j |
|  | P<sub>4</sub> | 7.47 ± 0.29hi | 4.40 ± 0.47h-j | 245.97 ± 5.47kl | 49.27 ± 1.97 | 216.72 ± 5.31jk | 59.50 ± 2.42g-i | 27.51 ± 0.69h-j | 17.90 ± 0.59f-j |
|  | P<sub>5</sub> | 9.67 ± 0.29b-d | 5.30 ± 0.3c-g | 341.50 ± 12.24a-d | 49.17 ± 1.66 | 329.08 ± 7.23a-c | 71.60 ± 0.81a-d | 41.84 ± 1.04a-c | 20.42 ± 0.46a-c |
| N<sub>5</sub> | P<sub>1</sub> | 8.87 ± 0.18d-f | 4.97 ± 0.15e-i | 313.64 ± 9.31c-g | 49.83 ± 1.43 | 287.39 ± 9.56fg | 66.77 ± 2.33c-f | 36.94 ± 1.12ef | 18.79 ± 0.75c-h |
|  | P<sub>2</sub> | 10.67 ± 0.18a | 6.00 ± 0.29a-d | 339.22 ± 7.97a-d | 50.63 ± 1.89 | 330.91 ± 6.53a-c | 71.90 ± 1.56a-c | 41.56 ± 0.82a-c | 19.85 ± 0.59a-e |
|  | P<sub>3</sub> | 6.43 ± 0.52jk | 4.23 ± 0.26ij | 248.70 ± 9.74kl | 47.70 ± 1.57 | 217.82 ± 6.47i-k | 58.00 ± 2.11g-i | 27.58 ± 0.76h-j | 16.93 ± 0.63h-j |
|  | P<sub>4</sub> | 6.93 ± 0.35ij | 4.27 ± 0.19h-j | 245.58 ± 1.62kl | 48.80 ± 2.71 | 214.29 ± 4.74jk | 58.40 ± 1.94g-i | 27.70 ± 0.27h-j | 18.31 ± 1.04e-j |
|  | P<sub>5</sub> | 10.00 ± 0.12ab | 5.83 ± 0.15a-c | 353.63 ± 12.04ab | 54.10 ± 2.23 | 337.58 ± 8.55a | 74.53 ± 2.31ab | 42.46 ± 1.23ab | 21.14 ± 0.87ab |
| LS |  | ** | * | ** | ns | * | * | * | * |

Values are means ± standard errors of three replications. Different letters within the column indicate statistically significant differences among the treatments according to LSD at $p \leq 0.05$ . Here, N<sub>1</sub>, N<sub>2</sub>, N<sub>3</sub>, N<sub>4</sub> and N<sub>5</sub> represent control (no fertilizer), 100, 110, 90 and 80 % of FRG'2018, respectively and P<sub>1</sub>, P<sub>2</sub>, P<sub>3</sub>, P<sub>4</sub> and P<sub>5</sub> indicate control (no PGR), GA<sub>3</sub>, NAA, 4-CPA and SA at 50 ppm, respectively. \* and \*\*: Significant at 5 % and 1 % level of probability, respectively, ns: Not significant, LS: Level of significance.

### Number of Leaflets

Significant variation (*p≤0.05*) in number of leaflets leaf^-1^ in tomato was noted due to the application of varied fertilizer doses and different types of PGRs (Table 3 and Table 4). Fertilizer treatment N_5_ resulted in maximum number of leaflets (8.58 leaf^-1^) which exhibited statistical harmony with N_3_ and N_4_ treatments. While, minimum number of leaflets leaf^-1^ was obtained in N_1_ treatment having statistical similarity with N_2_ and N_3_ treatments. Among the PGR treatments, maximum 10.19 leaflets leaf^-1^ was counted in plants under P_2_ treatment which was statistically dissonant from all other PGR treatments. On the reverse side, plants treated with P_3_ of PGRs had the least number of leaflets leaf^-1^ (6.38 leaflets leaf^-1^) followed by P_4_ treatment (6.91 leaflets leaf^-1^). Additionally, being significantly varied, interaction of nutrient and PGR revealed that plants under N_5_P_2_ had the highest leaflets (10.67 leaf^-1^) having statistical parity with N_2_P_2_, N_3_P_2_, N_4_P_2_ and N_5_P_5_ combinations. On the contrary, the least number of leaflets leaf^-1^ was noticed in N_1_P_3_ interaction (5.93 leaflets leaf^-1^) which had statistical similarity with N_2_P_3_, N_3_P_3_, N_3_P_4_ and N_5_P_3_ treatments (Table 4).

### Leaf Area

Remarkable variation (*p≤0.05*) among fertilizer doses, types of PGRs and their interactions in terms of single leaf area of tomato was observed (Table 3 and Table 4). Except N_1_ fertilization, all the fertilizer treatments had statistically similar leaf area where plants applied with N_3_ nutrient dose had the highest leaf area (306.13 cm^2^). Oppositely, N_1_ plants exhibited the lowest leaf area (286.33 cm^2^). Main effect of PGR showed that leaves of P_2_ treated plants possessed maximum area (337.66 cm^2^) being statistically identical with P_5_ treatment (327.75 cm^2^) while, minimum leaf area was determined in P_4_ treatment (247.89 cm^2^) followed by P_3_ treatment (270.77 cm^2^). Among the interactions, significantly the largest leaf was measured in N_3_P_2_ combination (357.33 cm^2^) which had statistical parity with N_2_P_5_, N_3_P_1_, N_4_P_2_, N_4_P_5_, N_5_P_2_ and N_5_P_5_ treatment combinations. While, the smallest leaf was attained in N_3_P_4_ treatment (225.03 cm^2^) being statistical consistent with N_1_P_4_, N_4_P_3_, N_4_P_4_, N_5_P_3_ and N_5_P_4_ interactions.

### Leaf SPAD value

Leaf relative greenness as per SPAD value of leaf didn’t differ significantly in case of main effect of fertilizer dose and interaction but variations were significant against PGR treatment (Table 3 and Table 4). Statistical superiority in leaf SPAD value was recorded in plants treated with P_2_ PGR (53.08) which exhibited statistical unity with P_5_ treatment (52.01). On the other hand, SPAD value of leaf was estimated in P_4_ treatment (47.87) having statistical parity with P_1_ (49.42) and P_3_ (48.15) treatments.

### Shoot Weight

Fertilizer and PGR application significantly (*p≤0.05*) influenced the fresh and dry weight of shoot of tomato under study (Table 3 and Table 4). Maximum fresh and dry weight of shoot was recorded in N_3_ treatment (283.94 g and 67.91 g, respectively) which was statistically alike with N_2_, N_4_ and N_5_ treatments while, minimum fresh and dry weight of shoot was resulted in N_1_ treatment (265.61 g and 62.34 g, respectively). Among the PGR treatments, P_2_ exhibited the highest fresh and dry weight of shoot (328.13 g and 73.20 g, respectively) which was statistically at par with P_5_ treatment for both fresh and dry weight of shoot (323.45 g and 72.49 g, respectively). Whereas, minimum shoot weight in fresh and dry basis was obtained in P_4_ treatment (216.82 g and 57.86 g, respectively) having statistical parity with P_3_ treatment for shoot dry weight (58.74 g). In addition, plants under N_3_P_2_ interaction had the highest fresh and dry weight of shoot (343.20 g and 76.17 g, respectively) which expressed statistical similarity with N_2_P_2_, N_2_P_5_, N_4_P_2_, N_4_P_5_, N_5_P_2_ and N_5_P_5_ combinations. Reversely, minimum fresh and dry weight of shoot was obtained in N_3_P_4_ (204.88 g) and N_1_P_4_ (53.47 g), respectively where other interactions of fertilizer with P_3_ and P_4_ had statistical similarity with those two combinations.

### Root Weight

Root fresh weight of non-fertilized (N_1_) tomato plants was measured statistically minimum (33.84 g) while all other fertilized plants had statistical similarity of which N_3_ had the maximum root fresh weight plant^-1^ (35.81 g). Root dry weight had non-statistical variation for fertilizer application (Table 3). In terms of PGR treatments, P_2_ exhibited the highest fresh and dry weight of root (41.47 g and 20.77 g, respectively) which was statistically at par with P_5_ treatment (41.10 g and 20.61 g, respectively). Whereas, minimum weight of root in fresh and dry basis was obtained in P_4_ treatment (27.39 g and 17.43 g, respectively) having statistical parity with P_3_ treatment (28.65 g and 17.79 g, respectively). Interaction of fertilizer and PGR exhibited that significantly the highest fresh and dry weight of root was registered in N_3_P_2_ (42.82 g) and N_4_P_2_ (21.49 g), respectively where N_2_P_2_, N_2_P_5_, N_4_P_2_, N_4_P_5_, N_5_P_2_ and N_5_P_5_ had statistical harmony for root fresh weight and N_1_P_2_, N_1_P_5_, N_2_P_2_, N_2_P_5_, N_3_P_1_, N_3_P_2_, N_3_P_5_, N_4_P_5_, N_5_P_2_ and N_5_P_5_ got statistical unity for root dry weight with the best combination. Contrarily, the lowest fresh and dry weight of root was obtained from N_3_P_4_ (26.12 g) and N_1_P_4_ (16.46 g) combinations, respectively (Table 4).

### Days Required to Flowering

Fertilizer application didn’t influence the number of days required to flowering in tomato; rather types of PGR significantly (*p≤0.05*) affected the floral induction date (Table 5). Due to fertilization, days required to flowering ranged from 47.33 days (N_3_) to 48.44 days (N_1_). Again, the earliest flowering within 44.02 days was found in P_4_ which expressed statistical unity with P_3_ treatment (44.27 days) while, delayed flowering was recorded in P_5_ treatment (51.35 days) having statistical affinity with P_1_ (50.05 days) and P_2_ (50.68 days) treatments. Moreover, nutrient-PGR interaction significantly influenced transplanting to floral induction duration in tomato where flowering occurred in the shortest possible time of 43.00 days in N_3_P_3_ combinations being statistically consistent with P_3_ and P_4_ interacted all fertilizer doses. On the other hand, N_4_P_5_ combination required maximum time (51.67 days) from transplanting to flowering in tomato which showed statistical similarity with the interaction treatments of all fertilizer doses and P_1_, P_2_ and P_5_ (Table 6).

**Table 5.**
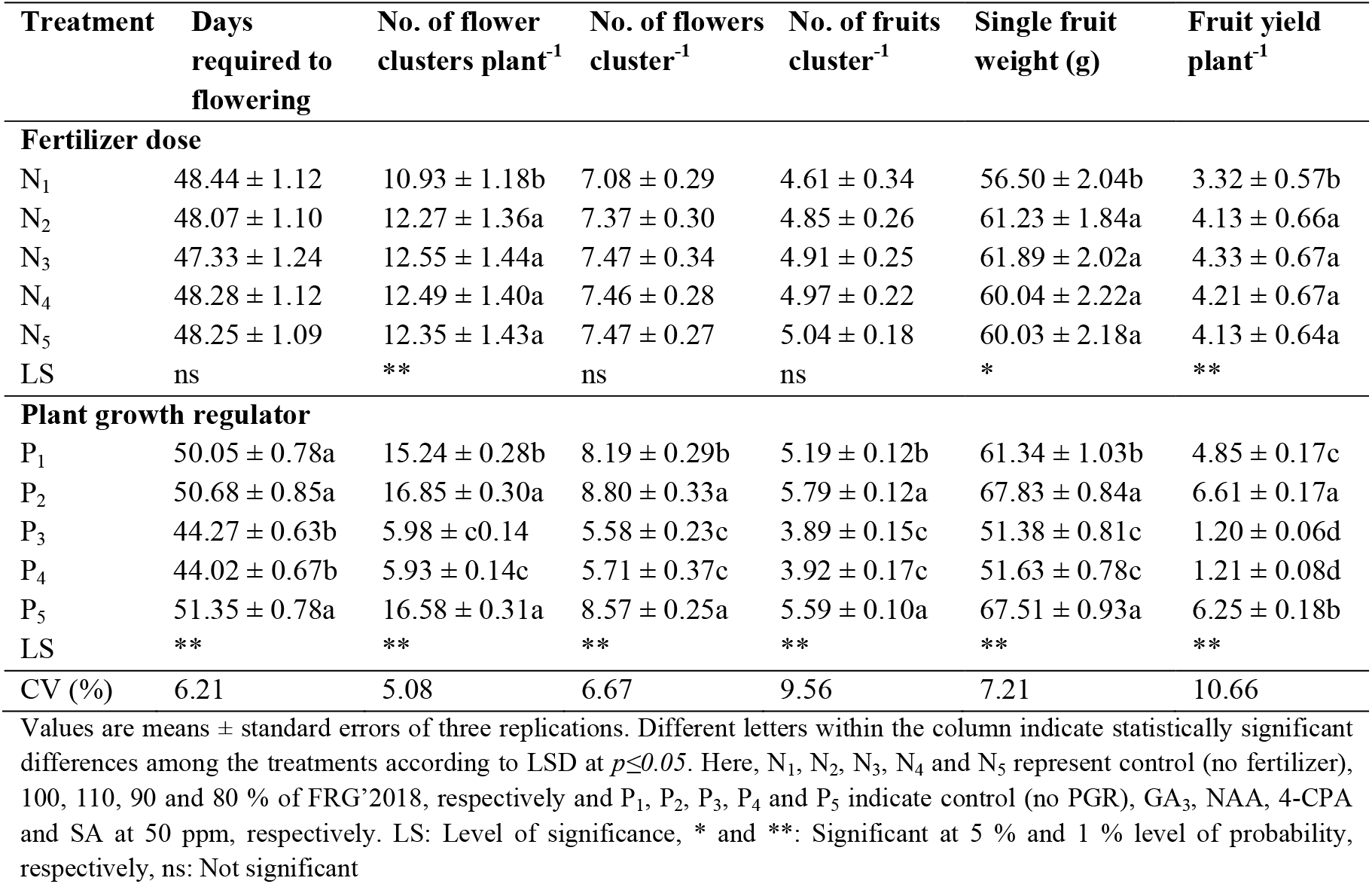
Reproductive behaviors and yield of tomato as influenced as influenced by the application fertilizers and plant growth regulators.

| Treatment | Days required to flowering | No. of flower clusters plant<sup>-1</sup> | No. of flowers cluster<sup>-1</sup> | No. of fruits cluster<sup>-1</sup> | Single fruit weight (g) | Fruit yield plant<sup>-1</sup> |
| --- | --- | --- | --- | --- | --- | --- |
| **Fertilizer dose** |  |  |  |  |  |  |
| N<sub>1</sub> | 48.44 ± 1.12 | 10.93 ± 1.18b | 7.08 ± 0.29 | 4.61 ± 0.34 | 56.50 ± 2.04b | 3.32 ± 0.57b |
| N<sub>2</sub> | 48.07 ± 1.10 | 12.27 ± 1.36a | 7.37 ± 0.30 | 4.85 ± 0.26 | 61.23 ± 1.84a | 4.13 ± 0.66a |
| N<sub>3</sub> | 47.33 ± 1.24 | 12.55 ± 1.44a | 7.47 ± 0.34 | 4.91 ± 0.25 | 61.89 ± 2.02a | 4.33 ± 0.67a |
| N<sub>4</sub> | 48.28 ± 1.12 | 12.49 ± 1.40a | 7.46 ± 0.28 | 4.97 ± 0.22 | 60.04 ± 2.22a | 4.21 ± 0.67a |
| N<sub>5</sub> | 48.25 ± 1.09 | 12.35 ± 1.43a | 7.47 ± 0.27 | 5.04 ± 0.18 | 60.03 ± 2.18a | 4.13 ± 0.64a |
| LS | ns | ** | ns | ns | * | ** |
| **Plant growth regulator** |  |  |  |  |  |  |
| P<sub>1</sub> | 50.05 ± 0.78a | 15.24 ± 0.28b | 8.19 ± 0.29b | 5.19 ± 0.12b | 61.34 ± 1.03b | 4.85 ± 0.17c |
| P<sub>2</sub> | 50.68 ± 0.85a | 16.85 ± 0.30a | 8.80 ± 0.33a | 5.79 ± 0.12a | 67.83 ± 0.84a | 6.61 ± 0.17a |
| P<sub>3</sub> | 44.27 ± 0.63b | 5.98 ± c0.14 | 5.58 ± 0.23c | 3.89 ± 0.15c | 51.38 ± 0.81c | 1.20 ± 0.06d |
| P<sub>4</sub> | 44.02 ± 0.67b | 5.93 ± 0.14c | 5.71 ± 0.37c | 3.92 ± 0.17c | 51.63 ± 0.78c | 1.21 ± 0.08d |
| P<sub>5</sub> | 51.35 ± 0.78a | 16.58 ± 0.31a | 8.57 ± 0.25a | 5.59 ± 0.10a | 67.51 ± 0.93a | 6.25 ± 0.18b |
| LS | ** | ** | ** | ** | ** | ** |
| CV (%) | 6.21 | 5.08 | 6.67 | 9.56 | 7.21 | 10.66 |

Values are means ± standard errors of three replications. Different letters within the column indicate statistically significant differences among the treatments according to LSD at $p \leq 0.05$ . Here, N<sub>1</sub>, N<sub>2</sub>, N<sub>3</sub>, N<sub>4</sub> and N<sub>5</sub> represent control (no fertilizer), 100, 110, 90 and 80 % of FRG'2018, respectively and P<sub>1</sub>, P<sub>2</sub>, P<sub>3</sub>, P<sub>4</sub> and P<sub>5</sub> indicate control (no PGR), GA<sub>3</sub>, NAA, 4-CPA and SA at 50 ppm, respectively. LS: Level of significance, \* and \*\*: Significant at 5 % and 1 % level of probability, respectively, ns: Not significant

**Table 6.** Fertilizer-PGR interaction influencing the reproductive behaviors and yield of tomato var. BARI Tomato-14.

| Treatment combination |  | Days required to flowering | No. of flower clusters plant<sup>-1</sup> | No. of flowers cluster<sup>-1</sup> | No. of fruits cluster<sup>-1</sup> | Single fruit weight (g) | Fruit yield (kg plant<sup>-1</sup>) |
| --- | --- | --- | --- | --- | --- | --- | --- |
| N<sub>1</sub> | P<sub>1</sub> | 49.00 ± 2.82a-c | 13.57 ± 0.59f | 8.27 ± 0.18ab | 5.27 ± 0.18a-c | 57.37 ± 2.35ef | 4.08 ± 0.14d |
|  | P<sub>2</sub> | 49.67 ± 2.53ab | 14.90 ± 0.67e | 8.77 ± 0.41a | 5.87 ± 0.33ab | 64.73 ± 1.97a-d | 5.63 ± 0.11b |
|  | P<sub>3</sub> | 43.00 ± 1.73d | 5.67 ± 0.20g | 4.83 ± 0.18e | 3.13 ± 0.18h | 48.33 ± 2.28g | 0.87 ± 0.11e |
|  | P<sub>4</sub> | 43.67 ± 1.86d | 5.67 ± 0.20g | 5.03 ± 0.52de | 3.27 ± 0.48gh | 48.90 ± 1.76g | 0.89 ± 0.09e |
|  | P<sub>5</sub> | 51.33 ± 2.23a | 14.87 ± 0.81e | 8.50 ± 0.17ab | 5.50 ± 0.17a-c | 63.17 ± 2.43a-e | 5.15 ± 0.29bc |
| N<sub>2</sub> | P<sub>1</sub> | 50.00 ± 2.08ab | 15.33 ± 0.20de | 7.87 ± 0.35b | 4.90 ± 0.38c-e | 61.50 ± 2.65c-e | 4.64 ± 0.47cd |
|  | P<sub>2</sub> | 50.67 ± 1.56a | 17.20 ± 0.49ab | 8.93 ± 0.26a | 5.93 ± 0.26a | 68.90 ± 3.69ab | 7.01 ± 0.38a |
|  | P<sub>3</sub> | 44.67 ± 1.59cd | 6.13 ± 0.30g | 5.67 ± 0.22cd | 3.77 ± 0.20f-h | 53.53 ± 0.92fg | 1.23 ± 0.06e |
|  | P<sub>4</sub> | 43.67 ± 1.59d | 6.10 ± 0.42g | 5.77 ± 0.47cd | 3.97 ± 0.39fg | 53.77 ± 1.55fg | 1.33 ± 0.24e |
|  | P<sub>5</sub> | 51.33 ± 2.03a | 16.57 ± 0.59a-c | 8.63 ± 0.19ab | 5.67 ± 0.18ab | 68.47 ± 2.77a-d | 6.43 ± 0.42a |
| N<sub>3</sub> | P<sub>1</sub> | 50.67 ± 1.76a | 16.10 ± 0.49b-d | 8.47 ± 0.41ab | 5.40 ± 0.35a-c | 64.30 ± 3.34a-e | 5.55 ± 0.19b |
|  | P<sub>2</sub> | 51.43 ± 2.5a | 17.50 ± 0.25a | 8.93 ± 0.35a | 5.80 ± 0.29ab | 68.47 ± 3.76a-d | 6.94 ± 0.47a |
|  | P<sub>3</sub> | 44.57 ± 1.6cd | 6.10 ± 0.49g | 5.77 ± 0.24cd | 3.97 ± 0.24fg | 53.87 ± 2.08fg | 1.29 ± 0.02e |
|  | P<sub>4</sub> | 44.43 ± 1.74cd | 5.90 ± 0.42g | 5.63 ± 0.35cd | 3.83 ± 0.35f-h | 53.13 ± 2.05fg | 1.23 ± 0.23e |
|  | P<sub>5</sub> | 51.10 ± 1.72a | 17.13 ± 0.30ab | 8.53 ± 0.35ab | 5.57 ± 0.33a-c | 69.70 ± 2.54a | 6.62 ± 0.27a |
| N<sub>4</sub> | P<sub>1</sub> | 50.10 ± 1.33ab | 15.67 ± 0.20c-e | 8.17 ± 0.24ab | 5.17 ± 0.24b-d | 61.40 ± 1.34de | 4.96 ± 0.11bc |
|  | P<sub>2</sub> | 51.30 ± 2.65a | 17.33 ± 0.20a | 8.90 ± 0.38a | 5.87 ± 0.35ab | 68.87 ± 2.35ab | 7.02 ± 0.61a |
|  | P<sub>3</sub> | 43.77 ± 1.13d | 6.23 ± 0.29g | 5.83 ± 0.24c | 4.17 ± 0.27ef | 50.53 ± 0.93fg | 1.32 ± 0.16e |
|  | P<sub>4</sub> | 44.57 ± 1.79cd | 6.00 ± 0.40g | 5.83 ± 0.26c | 4.13 ± 0.26ef | 50.83 ± 2.06fg | 1.26 ± 0.12e |
|  | P<sub>5</sub> | 51.67 ± 1.37a | 17.20 ± 0.49ab | 8.57 ± 0.26ab | 5.53 ± 0.23a-c | 68.57 ± 3.30a-c | 6.51 ± 0.27a |
| N<sub>5</sub> | P<sub>1</sub> | 50.47 ± 1.75a | 15.53 ± 0.39c-e | 8.20 ± 0.29ab | 5.20 ± 0.29a-c | 62.13 ± 2.94b-e | 5.00 ± 0.19bc |
|  | P<sub>2</sub> | 50.33 ± 1.37a | 17.33 ± 0.20a | 8.47 ± 0.26ab | 5.47 ± 0.26a-c | 68.17 ± 2.70a-d | 6.45 ± 0.28a |
|  | P<sub>3</sub> | 45.33 ± 1.7b-d | 5.77 ± 0.29g | 5.80 ± 0.23cd | 4.43 ± 0.23d-f | 50.63 ± 1.81fg | 1.29 ± 0.05e |
|  | P<sub>4</sub> | 43.77 ± 1.78d | 6.00 ± 0.17g | 6.27 ± 0.31c | 4.40 ± 0.31ef | 51.53 ± 1.88fg | 1.35 ± 0.06e |
|  | P<sub>5</sub> | 51.33 ± 2.68a | 17.13 ± 0.30ab | 8.60 ± 0.38ab | 5.70 ± 0.38ab | 67.67 ± 4.13a-d | 6.56 ± 0.14a |
| Level of significance |  | * | ** | * | * | * | ** |

Values are means ± standard errors of three replications. Different letters within the column indicate statistically significant differences among the treatments according to LSD at $p \leq 0.05$ . Here, N<sub>1</sub>, N<sub>2</sub>, N<sub>3</sub>, N<sub>4</sub> and N<sub>5</sub> represent control (no fertilizer), 100, 110, 90 and 80 % of FRG\*2018, respectively and P<sub>1</sub>, P<sub>2</sub>, P<sub>3</sub>, P<sub>4</sub> and P<sub>5</sub> indicate control (no PGR), GA<sub>3</sub>, NAA, 4-CPA and SA at 50 ppm, respectively. \* and \*\*: Significant at 5 % and 1 % level of probability, respectively.

### Number of Flower Clusters

Nutrient, PGR and their interaction significantly (*p≤0.05*) stimulated the attainment of flower clusters in tomato (Table 5 and Table 6). Number of flower clusters was counted minimum in non-fertilized (N_1_) plants (10.93 plant^-1^), while it was recorded maximum in N_3_ treatment (12.55 plant^-1^) being statistically at par with N_2_, N_4_ and N_5_ treatments. Among the PGR treatments, maximum number of flower clusters was noted in plants applied with P_2_ (16.85 plant^-1^) which had statistical unity with P_5_ treatment (16.58 plant^-1^), while minimum number of flower clusters was observed in P_4_ treatment (5.93 plant^-1^) being similar to that of P_3_ treatment (5.98 plant^-1^). Besides, interaction exhibited that plants under N_3_P_2_ treatment produced maximum number of flower clusters (17.53 plant^-1^) which had statistical harmony with N_2_P_2_, N_2_P_5_, N_3_P_5_, N_4_P_2_, N_4_P_5_, N_5_P_2_ and N_5_P_5_ treatment combinations. Whereas, N_1_P_3_ and N_1_P_4_ treated plants developed same minimum number of flower clusters (5.67 plant^-1^) being statistically consistent with all other interactions between fertilizer and P_2_ and P_4_ (Table 6).

### Number of Flowers and Fruits Cluster^-1^

Number of flowers and fruits cluster^-1^ didn’t vary significantly for fertilizer application but variation was significant in terms of PGR treatment and interaction of nutrient and PGR (Table 5 and Table 6). As a result of fertilization, number of flowers and fruits cluster^-1^ ranged from 7.08 to 7.47 and 4.61 to 5.04. Again, plants under P_2_ treatment had maximum 8.80 flowers and 5.79 fruits cluster^-1^ which showed statistical parity with P_5_ treatment (8.57 flowers and 5.59 fruits cluster^-1^). Oppositely, minimum number of flowers and fruits (5.58 and 3.89 cluster^-1^, respectively) was observed in P_3_ treatment having statistical harmony with P_4_ treatment. Once again, maximum 8.93 flowers and 5.93 fruits cluster^-1^ was counted in N_2_P_2_ and N_3_P_2_ interactions and N_3_P_2_ combination, respectively which exhibited statistical similarity with N_1_P_1_, N_1_P_2_, N_1_P_5_, N_2_P_5_, N_3_P_1_, N_4_P_2_, N_4_P_5_, N_5_P_1_, N_5_P_2_ and N_5_P_5_ interactions.

### Single Fruit Weight

Fertilizers and PGRs application resulted in statistical variation in single fruit weight of tomato (Table 5). Plants under control fertilizer treatment (no fertilizer) produced the lightest fruit (56.50 g), while rest other fertilizer treatments produced fruits having statistically identical weight of which N_3_ had heavier fruits (61.89 g). Among the PGR treatments, single tomato was weighed maximum in P_2_ treatment (67.83 g) which had statistical similarity with P_5_ PGR treated plants (67.51 g). On the other hand, tomatoes having minimum weight were harvested from P_3_ treated plants (51.38 g) being statistically at par with P_4_ Treatment (51.63 g). Furthermore, interaction treatment showed that plants under N_3_P_5_ treatment produced the heaviest tomato (69.70 g) having statistical parity with that of N_1_P_2_, N_1_P_5_, N_2_P_2_, N_2_P_5_, N_3_P_1_, N_3_P_2_, N_4_P_2_, N_4_P_5_, N_5_P_2_ and N_5_P_5_ combinations. All the nutrient treatments interacted with P_3_ and P_4_ PGRs exhibited statistical unity for producing tomatoes having minimum individual fruit weight where the lightest tomato was found in N_2_P_3_ interaction (48.33 g) (Table 6).

### Fruit Yield

As a result of variations in the vegetative and reproductive behaviors of tomato due to the application of fertilizers and PGRs, fruit yield of tomato also varied significantly (*p≤0.05*) in single effect as well as in interaction (Table 5 and Table 6). It was noticed that N_1_ fertilizer treatment had statistically minimum fruit yield (3.32 kg plant^-1^). Fertilizer doses like N_2_, N_3_, N_4_ and N_5_ had statistical harmony with respect to fruit yield where maximum yield was estimated in N_3_ treatment (4.33 kg plant^-1^). Besides, showing statistical disparity among all the PGR treatments maximum fruit yield was attained in P_2_ treatment (6.61 kg plant^-1^) followed by P_5_ treatment (6.25 kg plant^-1^) while minimum tomato yield was noted in P_3_ treatment (1.20 kg plant^-1^) having statistical similarity with P_4_ treatment (1.21 kg plant^-1^) and followed by P1 treatment (4.85 kg plant^-1^). Once again, interaction revealed that plants under N_4_P_2_ combination produced the highest yield (7.02 kg plant^-1^) and N_1_P_3_ combination had the lowest yield (0.87 kg plant^-1^). Treatment combinations like N_2_P_2_, N_2_P_5_, N_3_P_2_, N_3_P_5_, N_4_P_5_, N_5_P_2_ and N_5_P_5_ showed statistical parity with the best treatment, while N_1_P_4_, N_2_P_3_, N_2_P_4_, N_3_P_3_, N_3_P_4_, N_4_P_3_, N_4_P_4_, N_5_P_3_ and N_5_P_4_ interactions had statistical consistency with the worst yielding treatment combination in tomato under the study.

### Correlation Coefficient Analysis

Pearson’s correlation co-efficient indicated the interrelationships among the studied 29 variables including growth and yield attributing characters of tomato upon fertilizer and PGR treatment (Figure 7). It was noted that plant height, base diameter, number of branches and leaves plant^-1^ and canopy spread at 20 DAT (PH1, BD1, BN1, LN1 and CS1) exhibited very weak correlation (-0.09 to 0.52) to other vegetative and reproductive parameters indicating that initial plant growth hardly influenced the yield contributing attributes of tomato. Reproductive trait namely days required to flowering (DFL) was not affected much with the change in the plant growth attributes as there had low to moderate positive correlation with DFL to vegetative traits. SPD also had moderate positive correlation (0.46 to 0.66) with yield characters and low to moderate positive correlation with morphological growth features demonstrating that plant growth didn’t influence the leaf SPAD value and SPAD value didn’t influence the flowering and fruiting in tomato largely. Again, PH2, PH3, BD2, BD3, BN2, BN3, LN2, LN3, CS2, CS3, LLF, ITD, LFA, SFW, RFW, SDW and RDW expressed moderate to very strong positive correlation (0.67 to 0.95) with the yield and yield contributing variables namely FCN, FLC, FTC, FWT and FYP indicating that plant height, base diameter, number of branches and leaves plant^-1^, canopy spread at 45 and 75 DAT, leaflets leaf^-1^, internode length and single leaf area at full blossom as well as shoot and root fresh and dry weight had great impact on the flower clusters plant^-1^, flowers and fruits cluster^-1^, single fruit weight and fruit yield plant^-1^ in tomato upon the application of fertilizers and PGRs at vegetative growth stages.

**Fig. 7.**
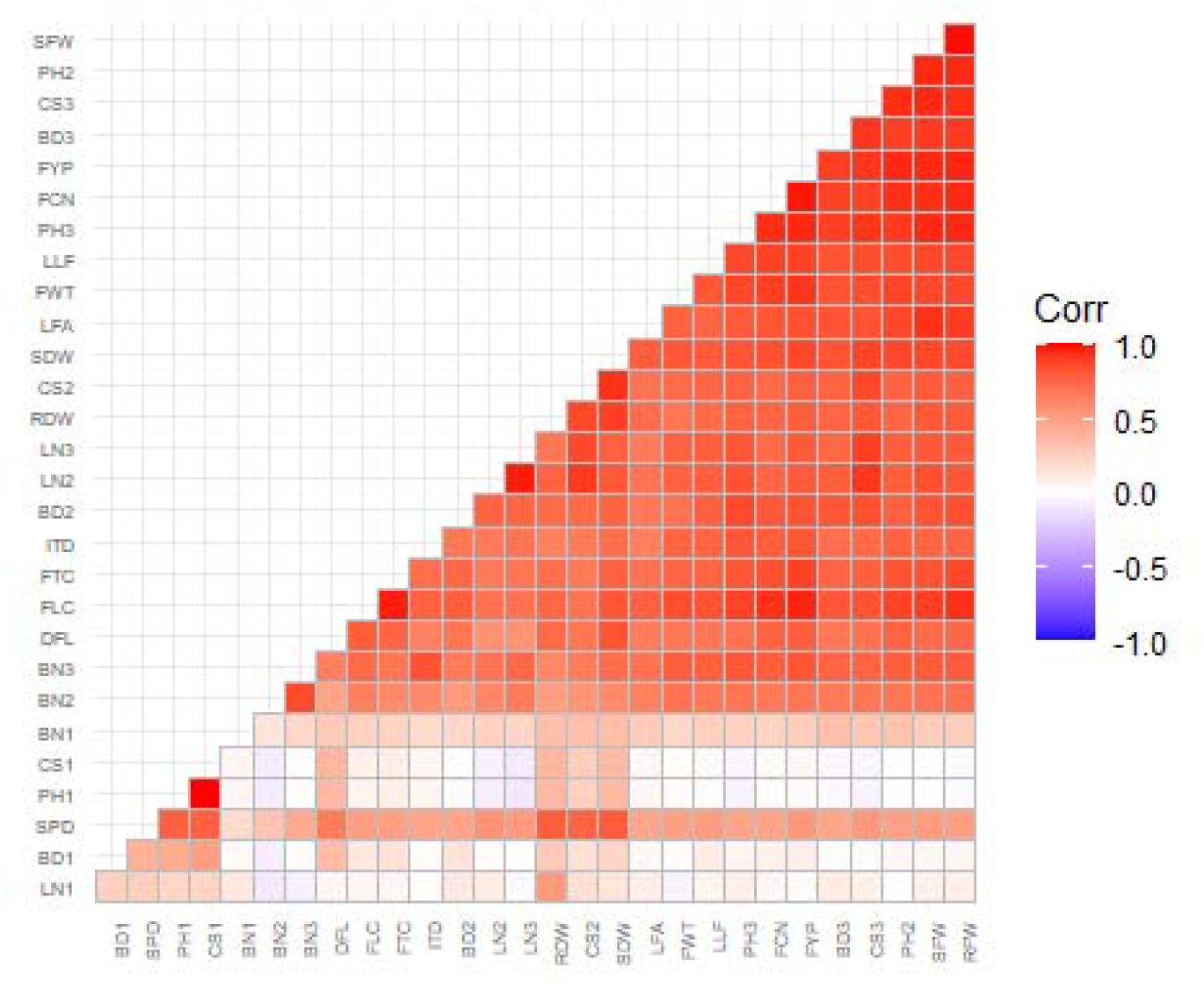
Correlation matrix of growth and yield related 29 variables of tomato. [PH, BD, BN, LN and CS represent plant height, base diameter, number of branches plant^-1^, number of leaves plant^-1^ and canopy spread plant^-1^, respectively and the adjacent digits 1, 2 and 3 indicate 20, 45 and 75 DAT, respectively; LLF, ITD, LFA and SPD allude number of leaflets leaf^-1^, internode length, single leaf area and leaf SPAD value, respectively; SFW, SDW, RFW and RDW indicate shoot and root fresh and dry weight, respectively; DFL, FCN, FLC, FTC, FWT and FYP represent days required to flowering, number of flower clusters plant^-1^, number of flowers cluster^-1^, number of fruits cluster^-1^, individual fruit weight and fruit yield plant^-1^, respectively]

### Principal Component Analysis

Principal component analysis (PCA) was employed to depict the relationship and impact of different fertilizer treatments and PGR types on growth and yield of tomato under high day and low night temperature condition in the late winter. The first four principal components, namely PC1 to PC4 (eigenvalues ≥ 1), account for 85% of the total variance in the data set of which PC1 (Dim1) and PC2 (Dim2) explained 65.4 % and 12 % of the total variation, respectively (8A). As seen in Figure 8, both Dim1 (PC1) and Dim2 (PC2) were positively correlated with plant height, base diameter, number of branches and leaves plant^-1^, canopy spread at 45 and 75 DAT (PH2, PH3, BD2, BD3, LN2, LN3, CS3), leaflets leaf^-1^, internode length and single leaf area at full blossom (LLF, ITD, LFA), shoot and root fresh weight (SFW, RFW), flower clusters plant^-1^ (FCN), flowers and fruits cluster^-1^ (FLC, FTC), single fruit weight (FWT) and fruit yield plant^-1^ (FYP) in tomato after the application of fertilizers and PGRs. These variables were, therefore, the most contributing factors for determining the best treatment of fertilizer and PGR.

**Fig. 8.**
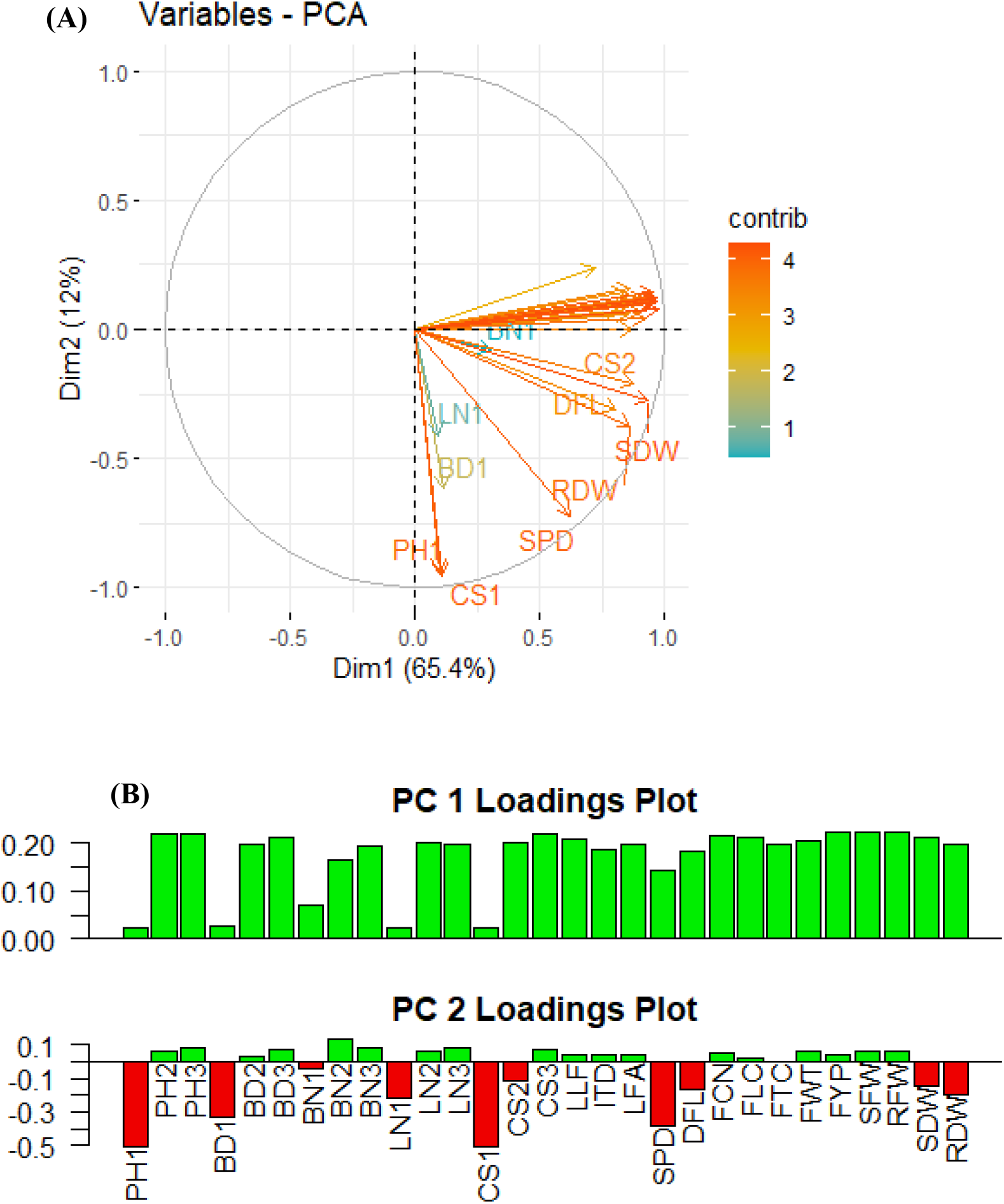
Principal component analysis (PCA) (A) and factor loadings for the first two principal components (Dim 1 and Dim 2) (B) of growth and yield attributes of tomato. [PH, BD, BN, LN and CS represent plant height, base diameter, number of branches plant^-1^, number of leaves plant^-1^ and canopy spread plant^-1^, respectively and the adjacent digits 1, 2 and 3 indicate 20, 45 and 75 DAT, respectively; LLF, ITD, LFA and SPD allude number of leaflets leaf^-1^, internode length, single leaf area and leaf SPAD value, respectively; SFW, SDW, RFW and RDW indicate shoot and root fresh and dry weight, respectively; DFL, FCN, FLC, FTC, FWT and FYP represent days required to flowering, number of flower clusters plant^-1^, number of flowers cluster^-1^, number of fruits cluster^-1^, individual fruit weight and fruit yield plant^-1^, respectively]

Again, the PCA-biplot exhibited that the entire fertilizer treatments outlays one-another without forming any distinct separate clusters expressing slight deviation of N_2_, N_3_, N_4_ and N_5_ from N_1_ in relation to the Dim1 and Dim2 which represents those fertilizers had little or no influence on the growth and yield of tomato under studied condition (Figure 9A). On the other hand, Figure 9B shows that the five PGR treatments including control are grouped into three different clusters namely cluster I (P_3_ and P_4_), cluster II (control) and cluster III (P_2_ and P_5_) where P2 is located at a distinct position in the positive quadrant with relation to Dim1 and Dim2. The treatment P_5_ overlaps P_2_ in the right quadrants (Q1 and Q2) exhibiting that there existed close statistical similarity having positive correlations among the plant growth and yield contributing characters in tomato. Meanwhile, P_3_ and P_4_ outlaying one-another also had a distinguished position at the left side in the PCA-biplot which demonstrate that these two PGRs had negative impact on tomato at the studied condition. Whereas, P_1_ obtained the central position in the PCA and distributed in all the quadrants having slight overlapping with P_2_ and P_5_ at the right side which represent that the treatment contributed positively for most of the parameters. Further, the longer vector length of the parameters could better represent PC1 and PC2, and the results indicated the relationship between parameters by confirming the angle between two vectors (0° < positively correlated < 90°; uncorrelated, 0°; 90° < negatively correlated < 180°). Therefore, it can be concluded that P_2_ and P_5_ treatments had significant influence on promoting tomato growth and yield at the fluctuating temperature condition in the late winter.

**Fig. 9.**
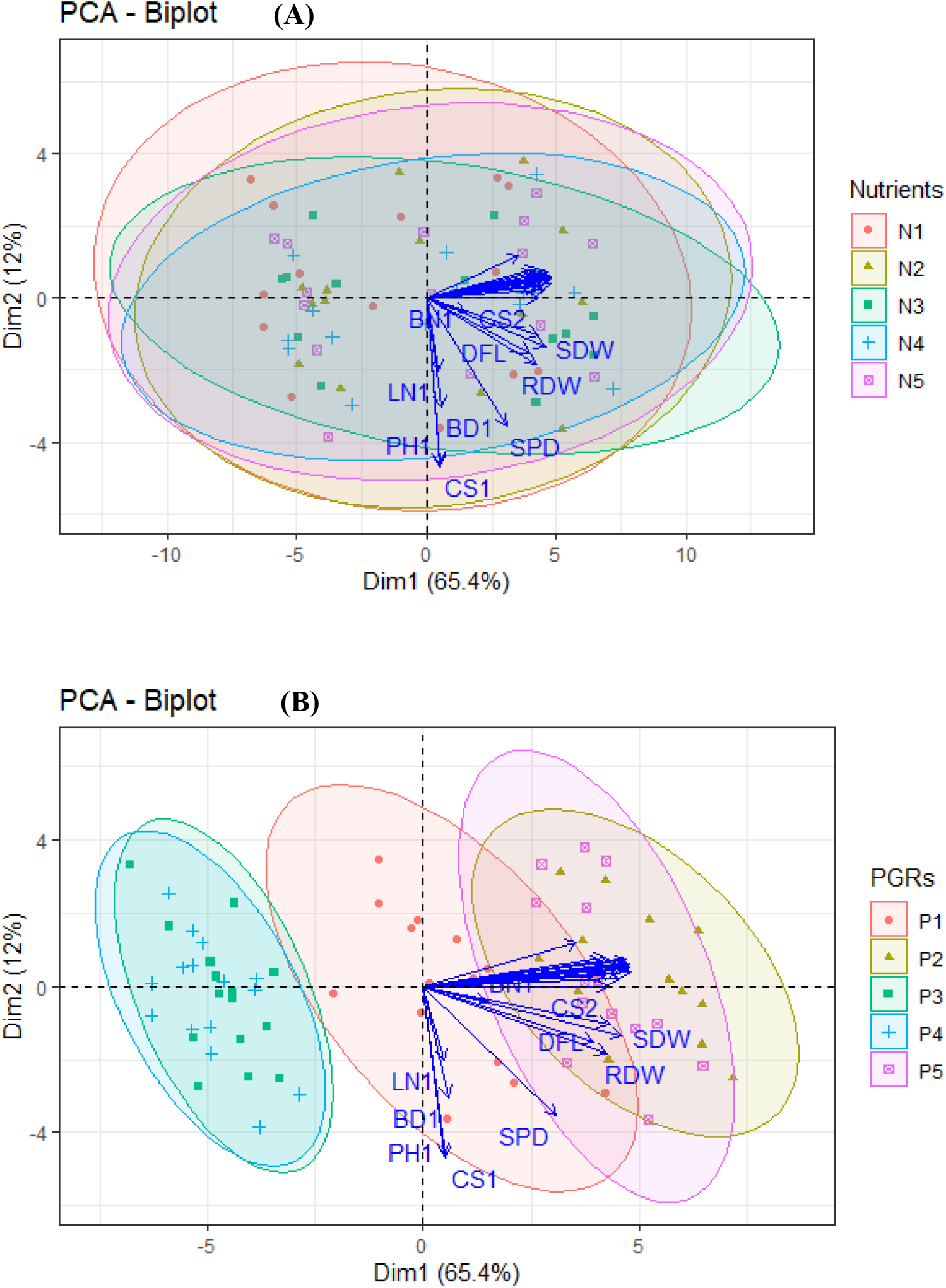
PCA-Biplot of the Nutrients and PGRs treatment. [PH, BD, BN, LN and CS represent plant height, base diameter, number of branches plant^-1^, number of leaves plant^-1^ and canopy spread plant^-1^, respectively and the adjacent digits 1, 2 and 3 indicate 20, 45 and 75 DAT, respectively; LLF, ITD, LFA and SPD allude number of leaflets leaf^-1^, internode length, single leaf area and leaf SPAD value, respectively; SFW, SDW, RFW and RDW indicate shoot and root fresh and dry weight, respectively; DFL, FCN, FLC, FTC, FWT and FYP represent days required to flowering, number of flower clusters plant^-1^, number of flowers cluster^-1^, number of fruits cluster^-1^, individual fruit weight and fruit yield plant^-1^, respectively]

## Discussion

Light, temperature and humidity are the major environmental factors influencing the plant physiological functions (Chiang et al., 2020; Mondaca et al., 2022), photosynthesis and hormonal balance (Driesen et al., 2020) and key active processes in plants life (Pant et al., 2021; Qaderi et al., 2019) including the transition from one development stage to the next (Chia & Lim, 2022; Dinu et al., 2022; Upadhyay et al., 2022). Besides environmental issues, edaphic elements also have remarkable influence on plant growth and development (Triantafyllidis et al., 2020) as soil nutrient availability is one of the absolute needs of plants (Gao et al., 2023a; Smith et al., 2022). But imbalances or deviations of these prime requirements from the optimum levels create stresses causing disruption in the physiological functioning, break down the hormonal balance and ultimately assert negative impact on crop yield and produce quality (Cannavo et al., 2022; Guo et al., 2022; Gupta et al., 2023).

In the present research, tomato var. *BARI Tomato-14*, a regular winter crop (Azad et al., 2020), has been grown in the late winter treating with different degrees of fertilization from 80 % to 110 % of recommendation (Ahmed et al., 2018). The plants were further foliarly applied with GA_3_, NAA, 4-CPA and SA @ 50 ppm at the vegetative stage. It was observed that growth and yield contributing traits didn’t vary statistically with the increment of fertilizer dose from 20 % less (80 %) to 10 % extra (110 %) of the recommended dose (100 %), though numerical enhancement in growth and yield of tomato was noticed with fertilizer increase. Rather, variations were only noted from control which means that fertilized plants performed better than non-fertilized plants but fertilizer increment or reduction to a certain level didn’t influence the plant responses statistically. On the other hand, the four types of PGRs exhibited notable statistical variations in morphological and reproductive responses of tomato under studied condition. Researchers addressed that nutrient application has significant positive impact on tomato growth, reproduction and yield (Gao et al., 2023b; Traoré et al., 2022) which are in resemblance with the present findings in terms of control versus fertilized plants and inconsistent in terms of the effect of the varied fertilizer doses. In the present observation, fluctuating as well as unstable temperature and humidity conditions during the late winter might have obstacle the efficient nutrient uptake by the tomato plants to response differently against varied levels of fertilization because plant performs negatively to any sorts of stresses. In addition, Kim et al. (2022) and Loudari et al. (2022) investigated that physiological functions, stomatal opening and hormonal regulations all get disrupted in imbalance weather conditions which restrict the nutrient uptake, plant growth and development as noted here with tomato.

Again, among the five PGR treatments including control, gibberellic acid (GA_3_) exhibited excellent vegetative and reproductive flourishment and salicylic acid (SA) had statistical resemblances with GA_3_ in most cases. Tomato plants under these two treatments had statistically maximum height, internode length, branching, leaves, canopy cover, fresh and dry biomass as well as flowers, fruits and finally tomato yield. Reversely, naphthalene acetic acid (NAA) and 4-chlorophenoxy acetic acid (4-CPA) treated plants showed retarded growth and development giving the ultimate lowest fruit yield. Plants receiving no PGR had mid-range responses. Tomato is a thermo-sensitive crop and plants are very much susceptible to changes in temperature, humidity and light and respond meticulously against stresses (Guo et al., 2022; Zheng et al., 2020). Again, plants’ responses to PGRs, especially when applied as spray, depend on environmental conditions like temperature, humidity, wind speed and light intensity (Cline & Bijl, 2002). Following foliar application of GA_3_ and SA, the growth characteristics of tomato plants were improved because these two PGRs had various effects on promoting cell division, cell enlargement/expansion, tissue differentiation, organ creation, vascular development, nutrient absorption, photosynthesisand biomass accumulation (Bagautdinova et al., 2022; Guo et al., 2022; Kaya et al., 2023; Shah et al., 2023). Again, GA_3_ and SA’s influence on stimulating the vegetative and reproductive behaviors of tomato under differential temperature and humidity conditions might be due to their ability to promote plants defense mechanisms against stresses (Ogugua et al., 2022). Ogugua et al. (2022) and Singh et al. (2019) noted that GA_3_, among different exogenous plant growth regulators, exhibited significant positive impact on superior growth and yield in tomato. In addition, Ali et al. (2022) and Guo et al. (2022) examined GA_3_ success in fluctuating summer temperature stress mitigation in tomato. GA_3_ is important for tomato production to boost yield and improve fruit quality under unfavorable climatic conditions of high temperature (Gelmesa et al., 2012). Besides, salicylic acid (SA) is one of the multifunctional hormones whose supplemental use in plants regulate physiological, biochemical, and photosynthetic pigments and molecular mechanisms in response to stressful conditions (Chen et al., 2023) and this ability of SA stimulate the tomato growth and yield in the present research under unstable temperature and humidity regimes. Endogenous hormone levels in plants, especially auxins, are also degraded with day-night temperature fluctuation and humidity alteration (Zheng et al., 2020). In the present investigation, shorter plant growth and subsequent lower yield in tomato under NAA and 4-CPA might be due to the environmental unrest that restrict tomato plants to response positively upon applications. Reduce plant height, branching, canopy led to the lower number of flowers and fruits in plants which ultimately inferior fruit yield with NAA and 4-CPA treatment though flourishing results are available with NAA and 4-CPA application (Ali et al., 2022; Jha et al., 2022). Again, flowering as well as fruit setting in tomato and other crops was promoted by GA_3_ at low concentration (Maboko & Du Plooy, 2015) as found in the present investigation too. Further, profound vegetative flourishment with higher number of branches and leaves as well as higher plant biomass in GA_3_ and SA treated plants accelerated the nutrient uptake from soil by plants (Emamverdian et al., 2020). Additionally, high fresh biomass and better canopy dimensions accounted for enhanced rate of photosynthesis and the latter process of accumulation and translocation of photosynthates to the sink resulting in the significant quantity of fruits having better quality in tomato during the late winter. Regulation in photosynthesis and source-sink translocation due to plant growth regulator application also noted in several studies (Katel et al., 2022; Shah et al., 2023). Consequently, PGRs application could suppress the fertilizer efficiency in the adverse situation and our correlation and PCA findings also substantiate these phenomena. Therefore, the use of GA_3_ and SA among studied PGRs revealed effective to minimize the fertilization up to 20% for enhancing the morphological and reproductive traits of tomato.

## Conclusion

In conclusion, the exogenous application of GA_3_ and SA @ 50 ppm remarkably influenced vegetative growth, yield-related reproductive traits and yield of tomato. Aside from this, fertilizer application 80-110 % of the recommended doses have no significant impact in compare to the PGRs that revealed that exogenous application of GA_3_ and SA (salicylic acid) could reduce 20% of chemical fertilization without compromising the tomato yield. Therefore, it can be suggested that application of GA_3_ and SA would be used as substitute of chemical fertilizer for enhancing the growth and yield of tomato even under adverse temperature and humidity differentiation in late winter in Bangladesh. Since this is an initial investigation on the use of PGRs as substitution of chemical fertilizer, further studies would be carried out with a wider range of PGRs concentrations focusing on the other abiotic stress conditions for comprehensive understanding of the interactive mode of action of the respective PGR in response to the specific growing stages of tomato.

## Acknowledgements

The authors are highly grateful to the Ministry of Science and Technology (MoST), Bangladesh for the financial support to continue this study under the project entitled “Plant growth regulators (PGRs) mediated approach in reducing chemical fertilizers for Tomato and Brinjal cultivation addressing adverse climate in Bangladesh” (Project ID: SRG-221180). The authors further would like to extend their sincere appreciation to the Researchers Supporting Project number (RSP2023R194), King Saud University, Riyadh, Saudi Arabia.

## Author contributions

J Hassan and J Gomasta conceived the idea of the study, design the experiment, conduct the study and wrote the manuscript. J Hassan, J Gomasta and H Sultana contributed in sample collection, preparation and laboratory analyses. J Hassan and J Gomasta analyze the data and made necessary interpretation. Y Ozaki, S Alamri, A T Alfagham and L A Al-Humaid reviewed and edited the manuscript for further improvement. All authors have read, edited the manuscript and approved it for submission.

